# Bi-level Graph Learning Unveils Prognosis-Relevant Tumor Microenvironment Patterns in Breast Multiplexed Digital Pathology

**DOI:** 10.1101/2024.04.22.590118

**Authors:** Zhenzhen Wang, Cesar A. Santa-Maria, Aleksander S. Popel, Jeremias Sulam

## Abstract

The tumor microenvironment is widely recognized for its central role in driving cancer progression and influencing prognostic outcomes. There have been increasing efforts dedicated to characterizing this complex and heterogeneous environment, including developing potential prognostic tools by leveraging modern deep learning methods. However, the identification of generalizable data-driven biomarkers has been limited, in part due to the inability to interpret the complex, black-box predictions made by these models. In this study, we introduce a data-driven yet interpretable approach for identifying patterns of cell organizations in the tumor microenvironment that are associated with patient prognoses. Our methodology relies on the construction of a bi-level graph model: (i) a cellular graph, which models the intricate tumor microenvironment, and (ii) a population graph that captures inter-patient similarities, given their respective cellular graphs, by means of a soft Weisfeiler-Lehman subtree kernel. This systematic integration of information across different scales enables us to identify patient subgroups exhibiting unique prognoses while unveiling tumor microenvironment patterns that characterize them. We demonstrate our approach in a cohort of breast cancer patients and show that the identified tumor microenvironment patterns result in a risk stratification system that provides new complementary information with respect to standard stratification systems. Our results, which are validated in two independent cohorts, allow for new insights into the prognostic implications of the breast tumor microenvironment. This methodology could be applied to other cancer types more generally, providing insights into the cellular patterns of organization associated with different outcomes.

## 1 Introduction

The tumor microenvironment (TME) is a complex ecosystem, comprising proliferating tumor cells, tumor stroma, immune cells, blood, and lymphatic vessels [1]. There is accumulating evidence underscoring the pivotal role of the TME in driving tumor progression [2], contributing to treatment resistance [3], and influencing patient prognosis [4]. Recent technological advancements in spatial multiplex proteomics, such as multiplexed immunohistochemistry (mIHC) [5], co-detection by indexing (CODEX) [6], multiplexed ion beam imaging (MIBI) [7], and imaging mass cytometry (IMC) [8], have enabled the simultaneous assessment of a wide spectrum of proteins in tissue specimens. These state-of-the-art technologies provide in-depth phenotypic profiling of hundreds of thousands of cells, allowing for a comprehensive exploration of the complexity and heterogeneity of TMEs at the single-cell level [9].

Increasing attention has been given to studying the TME at varying scales [10, 11], including the analysis of cell type compositions in various cancers [12–14], the quantification of spatial distances between pairs of cell phenotypes [15], and the characterization of cellular neighborhoods involving more than two cell types [16–18]. While these works have unveiled numerous unique patterns across a spectrum of cancers, associating them with clinical implications such as treatment response and patient prognoses remains an important and difficult challenge. In most cases, this question still relies on the formulation of explicit hypotheses regarding the relationship between the TME and diseases, grounded in domain expertise and prior knowledge [12, 19, 20]. However, this mostly hypothesis-driven approach may naturally constrain the exploration of novel relationships and patterns.

On the other hand, there is a growing interest in harnessing graph-based learning techniques to analyze the association between the TME and disease without the need for explicit hypotheses in a data-driven manner [21, 22]. Numerous recent studies have reported promising results by leveraging graph neural networks (GNNs) to model the TME and predict the presence [23, 24], grade [25, 26], stage [27], subtype [28, 29], and prognosis [30–34] of diverse types of cancers. Yet, the limited size, biological heterogeneity, and differences in staining and imaging protocols of clinical datasets constrain the quality of modern machine learning models by impeding their generalization capacity, particularly in cross-study scenarios that are often overlooked [35, 36]. Additionally, the interpretability of deep learning models, in general, remains a challenge [37–39]. Many of these studies do not discuss explanation in their works [25, 26, 28, 29, 32, 33], some employ gradient-based [23, 27, 34] and permutation-based [24] methods to generate an importance heatmap as the models’ explanation, and others conduct post-hoc analyses on learned graph features to obtain some clinical implications from their models [30, 31]. These explanations, although providing some insights into learned information, might not be clear, intuitive, or meaningful, especially for clinical experts and cancer researchers.

This study presents a novel method, named BiGraph, designed to discover characteristic TME patterns that are associated with good and bad prognosis, allowing for the stratification of patients. BiGraph addresses two key limitations of existing TME analysis methods: (i) it employs a data-driven approach, not relying on pre-defined, hand-crafted features typical of hypothesis-driven methods; and (ii) it is fully interpretable, allowing for the biological characterization of the found biomarkers, in contrast to the black-box nature of most graph deep learning methods. BiGraph entails the construction of two interconnected graphs: a patient-specific *cellular graph*, and a subsequent *population graph* given the patients’ characteristics captured by the former (see Fig. 1). The cellular graph models the cell spatial distribution of the TME for individual patients, capturing detailed information about the proximity, organization, and phenotypic information of cells extracted from tissue microarrays (TMA). On the other hand, the population graph captures similarities among all patients given their TME patterns at a macro, population level, with strong connections indicating high inter-patient similarities. The concept of graph-in-graph learning has been applied before in designing hierarchical GNNs [40, 41], with use cases in social networks [40], protein-protein interaction [42], and computational pathology [28]. BiGraph, on the other hand, introduces an efficient and interpretable method for connecting two hierarchies of graphs through a novel graph kernel function, referred to as the *Soft*-WL subtree kernel. The Soft-WL subtree kernel is a novel relaxation of the well-known Weisfeiler-Lehman (WL) subtree kernel [43], which involves the identification of heterogeneous TME patterns from patients’ cellular graphs and the comparison of the abundance of these patterns across patients (see Fig. 2). The combination of these graphs at different scales facilitates automatic and unsupervised patient risk stratification, where patients with similar TME patterns are clustered as *patient subgroups* through community detection methods, and prognoses of these subgroups are compared. Importantly, this risk stratification is also interpretable, as the features used to characterize each patient – histograms of TME patterns – are transparent and biologically meaningful. By comparing histograms within and outside a patient subgroup, the most characteristic TME patterns can be identified. In this way, the distinct survival outcomes observed among different patient subgroups provide valuable explicit hypotheses about the associations between TME patterns and patients’ prognoses, which can be systematically validated.

**Fig. 1:**
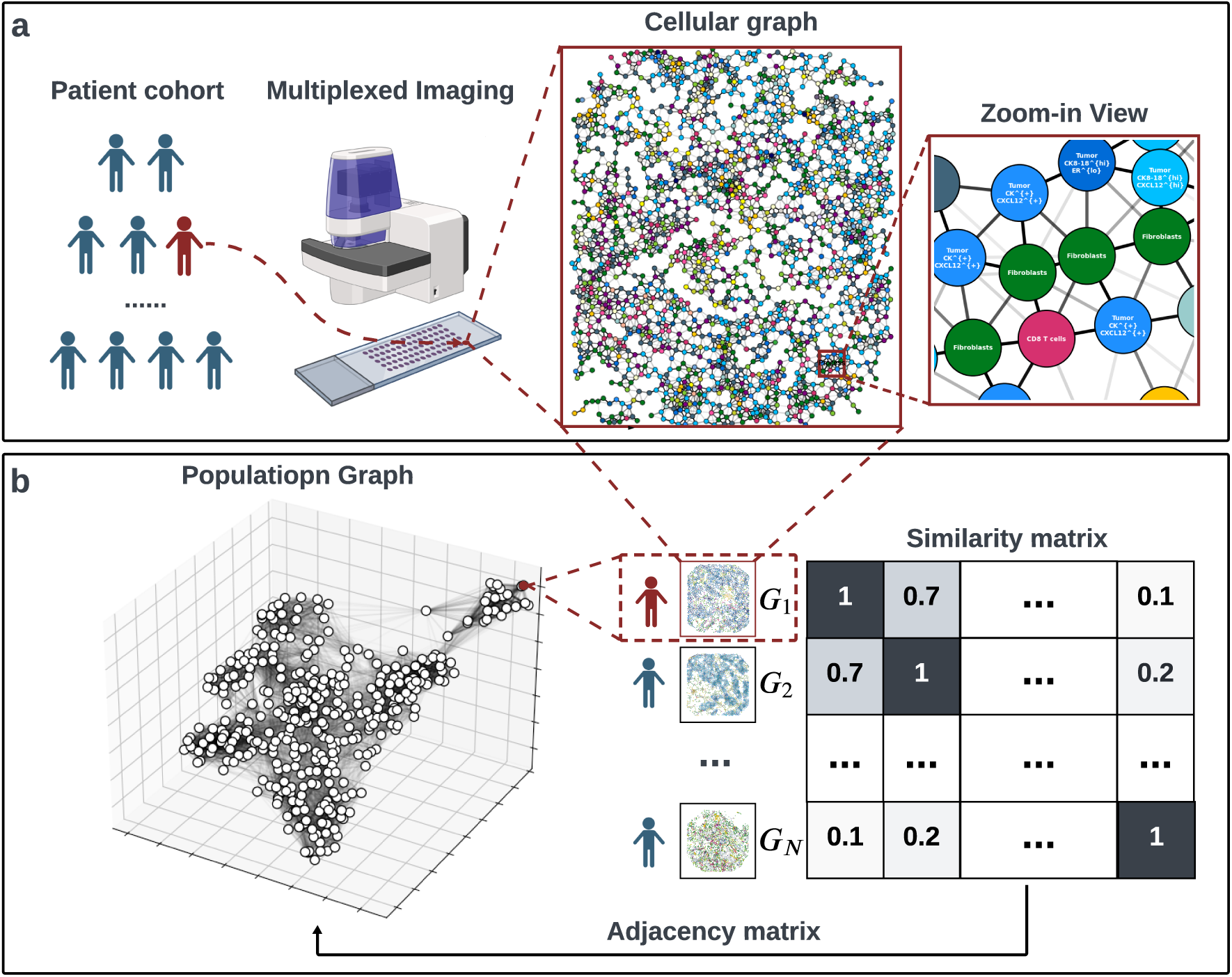
An overview of the BiGraph method. **(a)** The cellular graph is constructed to model patients’ tumor microenvironment (TME) based on their multiplexed images. Each node represents a cell, and cells are connected via edges with varying weights inversely correlated with inter-cellular distance. **(b)** The population graph takes the inter-patient similarity matrix as its adjacency matrix, where each node is a patient, and edge weight represents the inter-patient similarity.

**Fig. 2:**
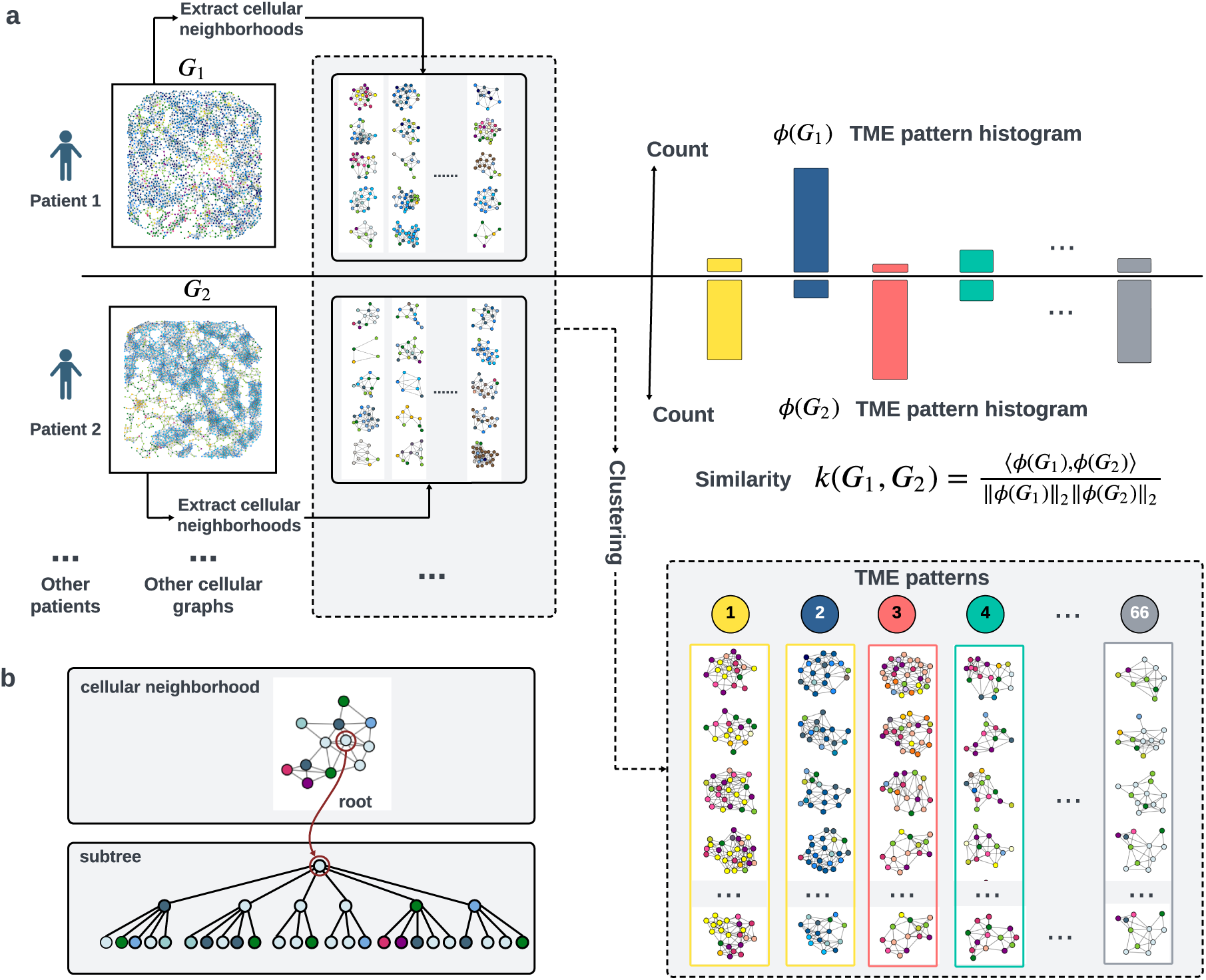
The Soft-WL subtree kernel. **(a)** Schematic diagram of the Soft-WL subtree kernel method. Numerous subtrees – each of which represents a cellular neighborhood – are extracted from cellular graphs. Similar subtrees are clustered together to form TME patterns. A TME pattern histogram is calculated for each cellular graph by counting the abundance of each TME pattern. The similarity between two cellular graphs is the normalized inner product of their TME pattern histograms. **(b)** The correspondence between a cellular neighborhood and a subtree. Given a cellular neighborhood with a designated root node, a subtree can be derived by iteratively identifying the neighborhoods of the root.

While the developed methodology is general, we focus our study on breast cancer. The analysis results demonstrate that BiGraph successfully provides a new risk stratification for breast cancer patients, which is complementary to the existing clinical subtyping system [44]; in other words, the found biomarkers provide insights that enhance the stratification provided by standard biomarkers of breast cancer, such as hormone receptor (HR) and human epidermal growth factor receptor 2 (HER2). Furthermore, the patient subgroups are characterized by specific, quantifiable TME patterns that lead to the discovery of prognosis-relevant TME patterns. To underscore the robustness of our findings, we validate them in both an inner validation set (comprising a randomly partitioned hold-out set from the same data source) as well as two external validation sets that have been independently collected and curated. These experiments showcase the generalizability of our results across diverse datasets, even in the presence of variations in antigen panels and cell phenotyping systems.

## 2 Results

In this section, we present our main results, starting with a high-level overview of the BiGraph method in Section 2.1. Section 2.2−2.5 demonstrates how BiGraph is applied to a breast cancer *discovery* cohort. Specifically, Section 2.2 describes the heterogenous TME patterns identified from cellular graphs. Section 2.3 explains how BiGraph measures inter-patient similarities. Section 2.4 details the patient stratification derived from the population graph, with a comparison with standard clinical attributes. Section 2.5 demonstrates the interpretability of BiGraph, where we interpret patient stratification results with characteristic TME patterns in each patient subgroup, followed by a systematic analysis of their prognostic values. In Section 2.6, we extensively validate the presented findings in inner- and external validation cohorts. Finally, Section 2.7 compares our method with alternative approaches.

### 2.1 BiGraph: Interconnected cellular graph and population graph capture multi-scale information

To tackle the identification of prognosis-relevant TME patterns, we develop an unsupervised bi-level graph learning method dubbed BiGraph. BiGraph is designed to be applicable to single-cell imaging data obtained through multiplexed imaging technologies. It takes as input tabular data containing the spatial coordinates and phenotypes of cells and outputs a risk stratification of patients as well as the prognosis-relevant patterns that characterize certain subgroups.

The primary innovation of BiGraph relies on exploiting relations across levels of graphs, namely, a cellular graph and a population graph, by means of a graph kernel method. The cellular graph models the TME of each individual patient, with each node in it corresponding to a cell and characterized by its spatial coordinates and phenotype label. Nodes (i.e., cells) are connected through edges representing inter-cellular interactions (see Fig. 1.a). Unlike conventional approaches that employ fixed distance thresholds to determine cell connectivity [15, 16, 30], we construct a complete cellular graph where all possible pairs of cells are potentially connected, with the strength of interaction decreasing as the distance between cells increases (see Section 5.2).

On the other hand, inspired by the PhenoGraph algorithm [45] that encodes phenotypic similarities of cells in a graph, our population graph models inter-patient similarities based on their TME patterns. Each node of the population graph represents a patient, and every patient is connected to potentially all other patients through edges with varying weights (see Fig. 1.b, Section 5.4). Each edge weight represents the similarity between the corresponding two patients, each characterized by their specific cellular graph. This similarity between patient’s cellular graphs is measured by a novel relaxation of the popular Weisfeiler-Lehman subtree kernel [43], namely the soft-WL subtree kernel. Such a kernel generates numerous *subtrees* from each cellular graph, each rooted in a single cell. A *feature embedding* is calculated for each subtree through an iterative graph convolution process. The subtree feature embedding encodes the unique composition and spatial organization of cells within a local cellular neighborhood surrounding the subtree root (see Section 5.3.1). Subtrees with similar embeddings are clustered to form TME *patterns* (see Section 5.3.2), and inter-patient similarity is established by comparing the abundance of TME patterns across patients (see Section 5.3.3). These notions of similarities between patients allow us to obtain clusters or sub-communities of patients. We employ the popular Louvain method [46] on the population graph to detect these communities, each representing a patient subgroup with common TME patterns (see Section 5.4). Importantly, the entire process is unsupervised: no information about the survival of patients is used to obtain the population graph. Survival analysis is subsequently applied to each patient subgroup to assess their relative risks. Furthermore, to better understand the distinct survival outcomes of these patient subgroups, the most characteristic TME patterns in each subgroup are identified, and their prognostic impacts are investigated (see Section 5.6).

To center our study of BiGraph in the context of breast cancer, we analyze a patient cohort curated by Danenberg et al. [16] with 693 breast cancer patients with heterogeneous clinical characteristics (Fig. 3.d), each accompanied by their 37-dimensional imaging mass cytometry (IMC) images of tissue microarray cores (TMA) and clinical data. This publicly available dataset provides well-established cell segmentation and phenotyping results, delineating 32 major cell phenotypes, as illustrated in Extended Data Fig. 1.a. From this dataset, 10 patients with no clinical data and 104 patients with less than 500 cells identified in their corresponding TMA images are excluded. The remaining 579 patients are randomly partitioned into two subsets containing 379 patients and 200 patients, respectively.

**Fig. 3:**
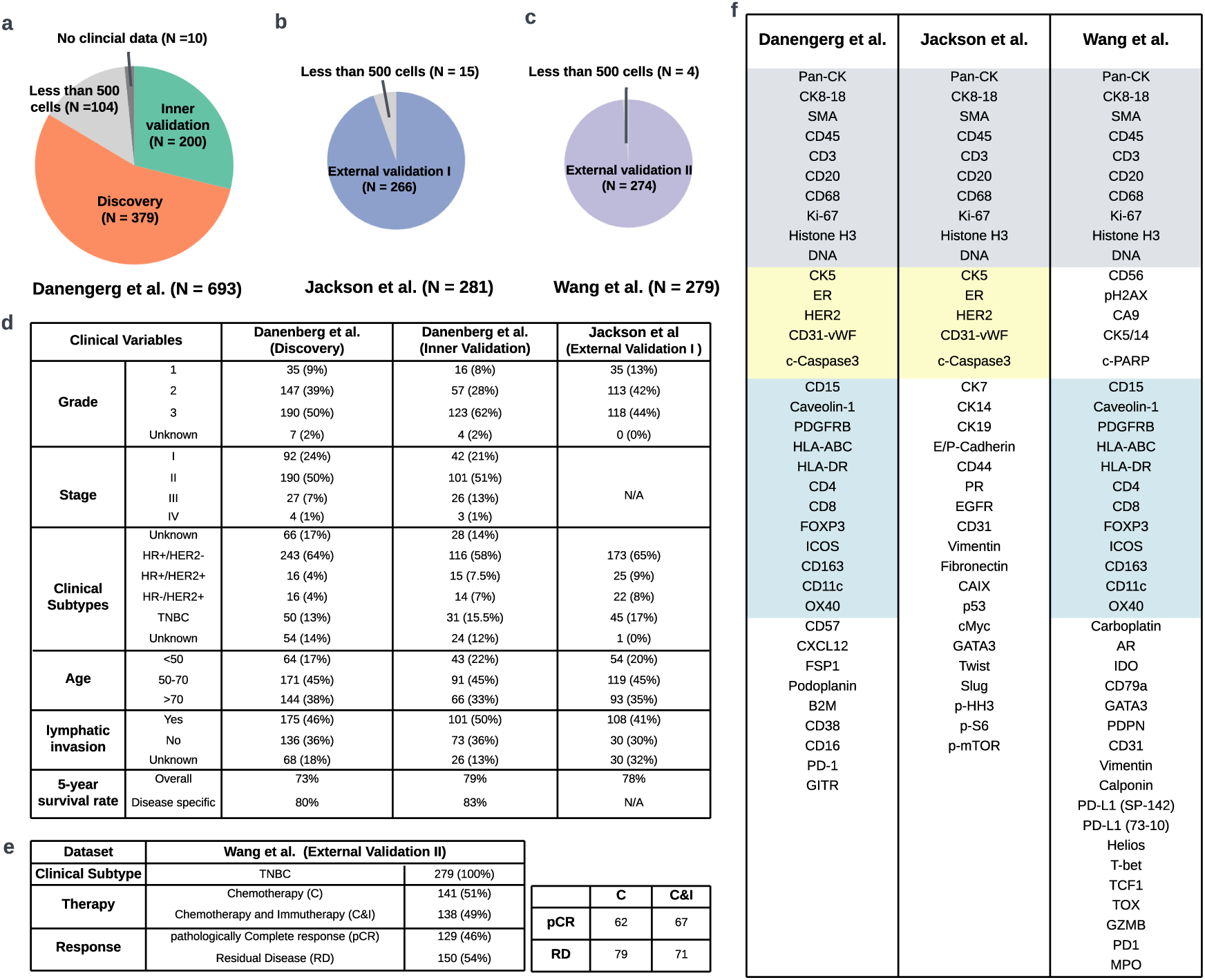
An overview of datasets. **(a)** Composition and partition for the 693 patients curated by Danenberg et al [16]. **(b)** Composition and partition for the 281 patients curated by Jackson et al (External validation set-1) [17]. **(c)** Composition and partition for the 281 patients curated by Wang et al (External validation set-2) [47]. **(c)** Clinical information of the discovery, inner validation, and external validation set-1. **(d)** Clinical information of the discovery, inner validation, and external validation set I. **(e)** Clinical information of the external validation set-2. **(d)** The antigens used in the datasets curated by Danenberg et al. [16], Jackson et al. [17], and Wang et al. [47]. Gray-shaded regions indicate the shared antigens by all three studies; yellow-shaded regions indicate the shared antigens by Danenberg et al. and Jackson et al.; blue-shaded regions indicate the shared antigens by Danenberg et al. and Wang et al.

We refer the larger subset with 379 patients to the *discovery set* and develop our BiGraph method based on it. The smaller subset with 200 patients is held out for validation purposes, referred to as the *inner validation set* (see Fig. 3.a). An external breast cancer patient cohort, independently curated by Jackson et al [17], is used as *external validation set-1*. This dataset contains 281 patients with heterogeneous clinical characteristics (Fig. 3.d), each accompanied by their 35-dimensional IMC images of TMAs and clinical data. The cell segmentation and phenotyping results are also publicly available (see Extended Data Fig. 1.b). Following the same patient exclusion criteria, 266 patients with more than 500 cells are included in the study (see Fig. 3.b). An additional external validation cohort, independently curated by Wang et al [47], is used as *external validation set-2*. This dataset comprises 279 early triple-negative breast cancer (TNBC) patients who participated in a randomized pre-surgical neoadjuvant immunotherapy clinical trial (NCT02620280). Although long-term follow-up data is not available, a binary label – pathological complete response (pCR), defined as the absence of invasive cancer cells in post-treatment tissues, or residual disease (RD) – is provided for each patient as a surrogate for prognosis. Tumor samples were collected at three distinct time points: before treatment, early on-treatment, and post-treatment. A total of 43 proteins are profiled in situ with the IMC technique. We analyze the pre-treatment and on-treatment samples to investigate the predictive value of TME patterns in response to therapy. As before, the cell segmentation and phenotyping results are also publicly available (see Extended Data Fig. 1.c). Four patients with less than 500 visible cells are excluded. Noticeably, the two external validation sets both employ a distinct antigen panel and phenotyping system from the discovery set – which is a common challenge in current biomarker discovery research – making this validation task challenging (see Fig. 3.f). Fortunately, about 43% (15/35) of the antigens used in the external validation set-1 and 51% (22/43) of the antigens used in the external validation-2 are shared with that in the discovery set, as shown in the shaded regions of Fig. 3.f. We use the shared antigens to match these two cell phenotyping systems and then conduct validation, as we detail in Section 5.7.1. For all the datasets, we use the processed data (i.e., cell segmentation and phenotyping results) instead of the raw IMC images as input to develop the BiGraph model.

### 2.2 Cellular graph analysis reveals heterogeneous TME patterns

The discovery set encompasses 379 breast cancer patients with their corresponding TMA images. For each patient, a cellular graph is constructed to model the spatial distribution of different types of cells within the TME. The Soft-WL subtree kernel generates numerous subtrees rooted at every single cell from each cellular graph. Each rooted subtree has a 32-dimensional feature embedding derived from an iterative graph convolution process, which encodes the composition of distinct cell phenotypes and their spatial arrangement in a local cellular neighborhood surrounding the subtree root (see Section 5.3.1).

The feature embeddings of subtrees allow us to cluster them, in an unsupervised fashion (see Section 5.3.2), culminating in a total of 66 clusters. Each of these clusters is considered a TME pattern, with unique composition and spatial organizations of cells. The proportion of each of them in every patient is calculated, revealing a high inter-patient heterogeneity of the prevalence (Fig. 4.a). A *signature* is assigned to each TME pattern to describe its unique composition and spatial organization of cells, which is formally defined as the average feature embedding of all the subtrees belonging to the designated TME pattern. The signature map reveals that the TME patterns exhibit heterogeneous features, and many of these involve intricate interactions among multiple cell phenotypes (Fig. 4.b). Based on the predominant cell phenotype, the TME patterns are grouped into four main categories: *tumor niches* that are mostly composed of tumor cells; *immune niches* with immune cells dominating; *stromal niches* that consist of almost all stromal cells; and *interface niches*, where tumor, immune, and stromal cells are interacting (Fig. 4.b). The vast majority of patients (i.e., n=283, 75%) have all these four categories in their cellular graphs, while their relative abundance varies across patients (Fig. 4.c).

**Fig. 4:**
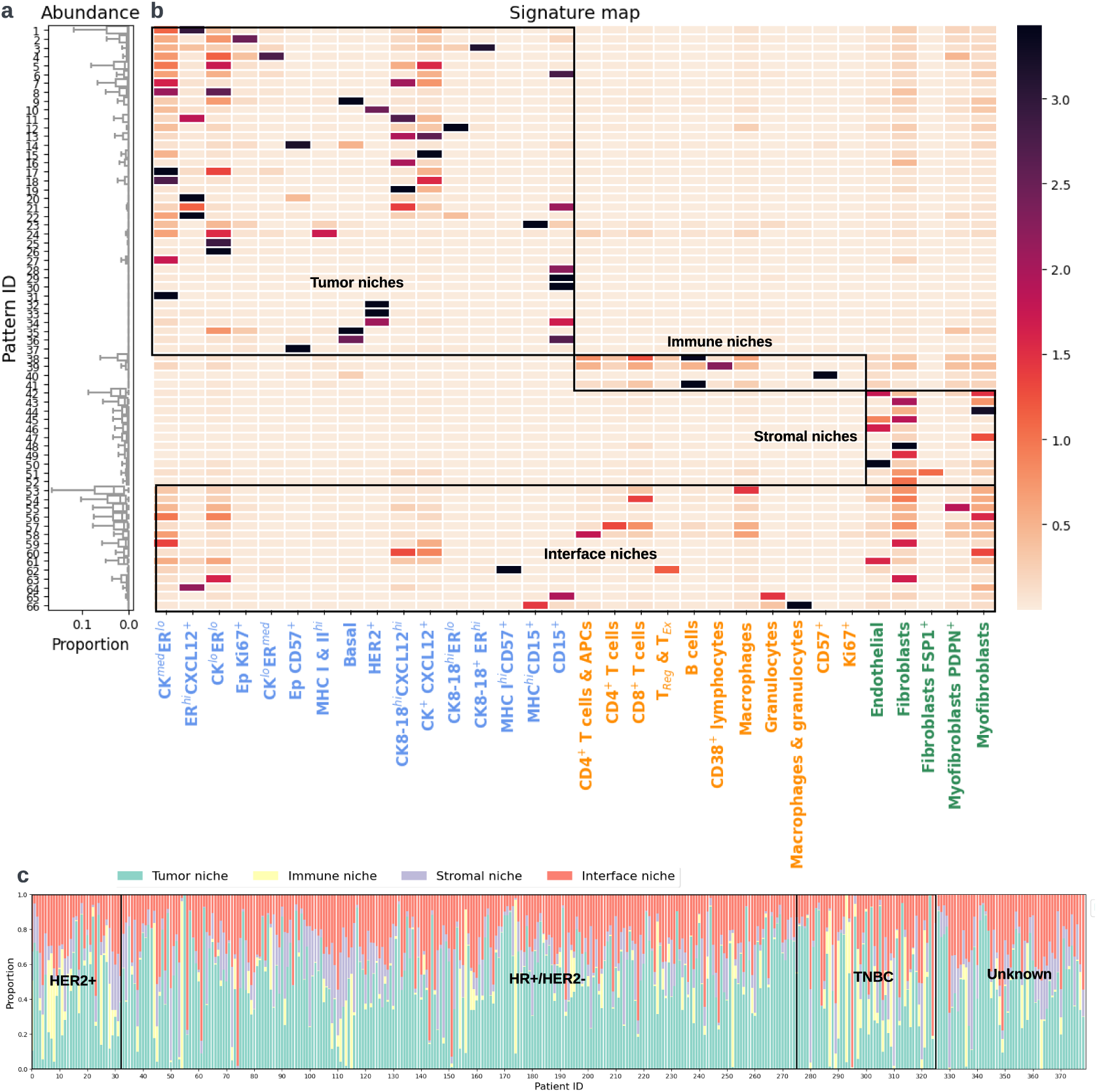
Abundance and Signature map of TME patterns. **(a)** Boxplot of the proportion of each TME pattern in each patient across the discovery set. **(b)** Signature map of these TME patterns; x-axis: cell phenotype; y-axis: pattern ID. x-axis text color indicates the cell type category: blue: tumor cells; orange: immune cells; green: stromal cells. Based on the dominated cell type, the 66 TME patterns are roughly classified into four classes: tumor niche, immune niche, stromal niche, and interface niche, as the bounding boxes indicate. **(c)** The proportion of immune niche, stromal niche, and interface niche in all patients.

### 2.3 Abundance of TME patterns provides inter-patient similarity measurement

Given the TME patterns extracted from each patient, we need a metric that precisely quantifies their similarities based on these patterns. Kernel functions are a natural choice. Specifically, the Soft-WL subtree kernel quantifies inter-patient similarity by comparing the abundance of these patterns in their respective TMEs. The similarity score between any pair of patients is determined by a normalized inner product of their histograms, describing the proportion of the 66 TME patterns (see Section 5.3.3, Equation (5)). The similarity score for any patient pair falls within the range [0, 1], with two identical graphs achieving a maximum similarity score of 1.

To illustrate the computation of inter-patient similarity, Fig. 5.a presents a randomly selected “template patient” from the discovery set (for illustration purposes) and depicts its cellular graph.

**Fig. 5:**
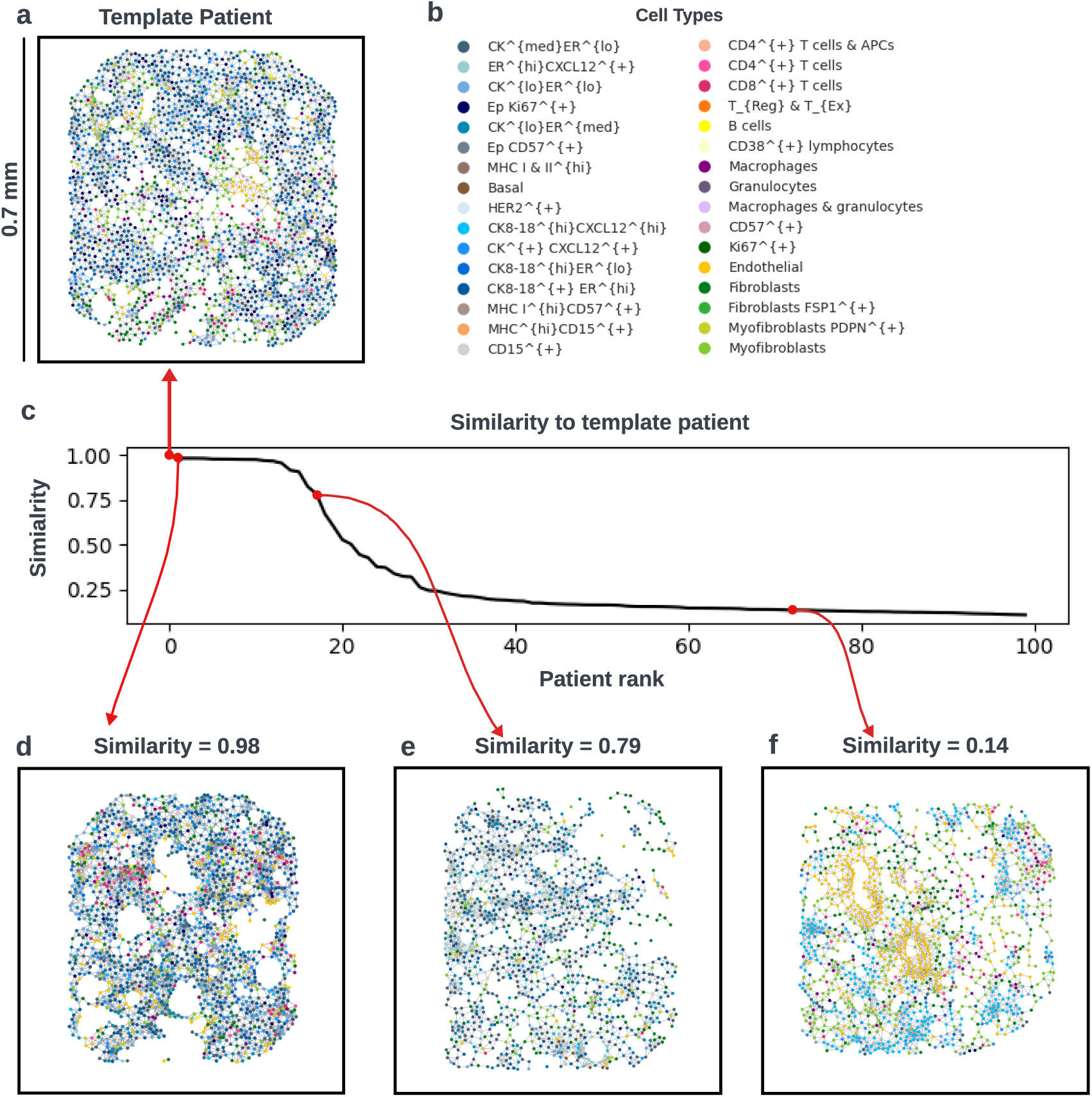
Demonstration of the inter-patient similarities measured by graph kernel methods. **(a)** The cellular graph of a selected patient, referred to as the “template patient”. **(b)** Colors of the 32 cell phenotype. **(c)** The similarity scores of other patients compared to the template patient (x-axis: patient ranking; y-axis: similarity score). Patients are sorted based on their similarity score, and only the top 100 patients are shown. **(d)-(f)** Cellular graphs of patients with varying similarity scores to the template patient.

The similarities between this template patient and all other patients are calculated and visualized, with the top 100 patients’ similarities shown in Fig. 5.b. To provide a more in-depth understanding of such comparison, we highlight three representative patients with varying degrees of similarity (see Fig. 5.d-f). The cellular graph of this template patient contains diverse tumor cells, with the predominant presence of cytokeratin-positive cells, including CK8/18^+^ER^high^ cells, CK8/18^high^CXCL12^high^ cells, CK^+^CXCL12^+^ cells, and CK^med^ER^low^ cells. Proliferative (i.e. Ki67^+^) tumor cells are also scattered throughout. Stromal, such as fibroblasts, myofibroblasts, and endothelial, intermingle with tumor cells, and a clear tumor-stromal boundary is difficult to discern. Immune cells, including CD4^+^ T cells, CD8^+^ T cells, and macrophages, are relatively rare and dispersed. Note how the patient with the highest similarity (similarity score of 0.98) exhibits nearly identical patterns to the template patient (see Fig. 5.d). Another patient with a similarity score of 0.79 (see Fig. 5.e) also features a substantial number of interactions among various cytokeratin-positive cells, including CK8/18^+^ER^high^ cells, CK8/18^high^CXCL12^high^ cells, and CK^+^CXCL12^+^ cells. However, distinct from the template patient, this patient includes many CK^low^ER^med^ cells and ER^high^CXCL12^+^ cells, resulting in the emergence of new patterns. Additionally, this patient exhibits a relatively immune-cold TME, with only a few immune cells infiltrating the tumor cells. In contrast, the last example patient demonstrates a substantially lower similarity score of 0.14, indicating significantly different patterns (Fig. 5.f). This patient presents with a vascular tumor, characterized by an abundance of endothelial cells. The tumor cells are relatively homotypic, dominated by CK8/18^high^CXCL12^high^ cells. This diversity in TME patterns illustrates the capability of the Soft-WL subtree kernel to discern nuanced differences among patients’ TMEs.

### 2.4 Population graph analysis provides data-driven risk stratification

Once we have a precise metric to quantify the similarity between any pair of patients, we proceed to construct a population graph that captures the inter-patient similarities across the entire set. In this graph, each node corresponds to an individual patient, and edges connecting nodes are assigned weights representing their respective similarity scores (see Section 5.4). A community detection algorithm is employed to reveal patient subgroups in an unsupervised manner without utilizing the patients’ survival information (see Section 5.4). The survival outcomes of these patient subgroups are analyzed and compared with the standard clinical subtyping system. We elaborate on these results in the following subsections.

#### 2.4.1 Patient subgroups exhibit significantly distinct survival outcomes

To identify patient subgroups characterized by similar TME patterns, we employ the widely used Louvain community detection algorithm [46] on the population graph. This process yields seven distinct communities, each representing a subgroup of patients exhibiting high intra-group similarities (see Section 5.4). These identified patient subgroups are denoted as S1 to S7. Notably, the nodes in the population graph lack spatial coordinates; hence, for visualization purposes, the Fruchterman-Reingold force-directed algorithm [48] is employed to depict a 3D representation of the population graph. This representation and the results of community detection (i.e., the resulting patient subgroups) are depicted in Fig. 6.a.

**Fig. 6:**
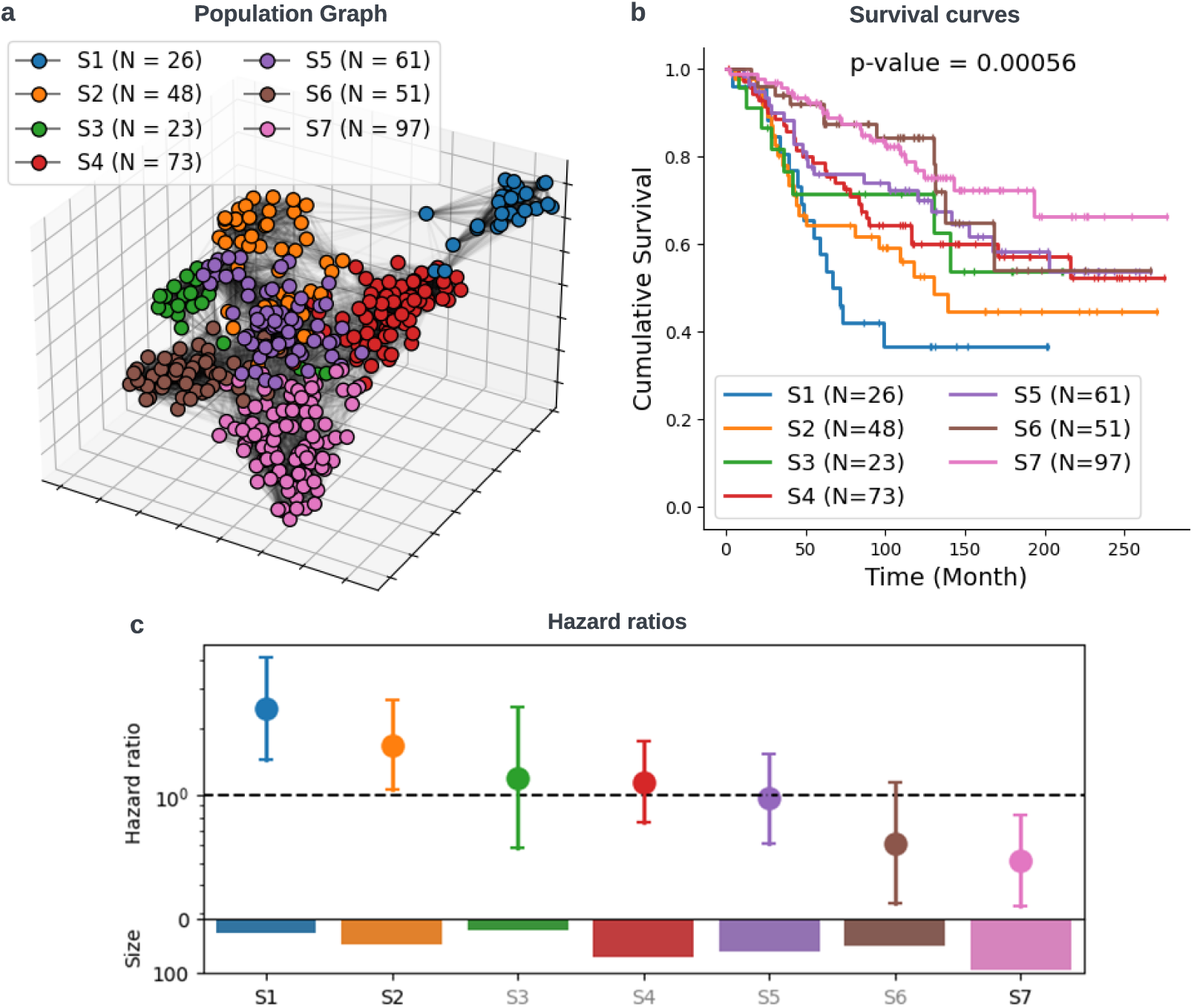
Population graph facilitating risk stratification. **(a)** Three-dimensional visualization of the population graph and the results of community detection. **b** Kaplan-Meier (K-M) survival plots depicting seven distinct patient subgroups. A multivariate log-rank test is conducted to compare the survival curves, with the resulting p-value indicated in the title. **c** Relative hazard ratios (with 95% confidence intervals) of the seven patient subgroups, estimated using a Cox proportional hazard model. Text color (grey vs. black) distinguishes statistically significant and non-significant associations (black: statistically significant). The size (i.e., number of patients) of each subgroup is presented in the corresponding barplot.

Disease-specific survival functions of the seven patient subgroups are estimated using the KaplanMeier (K-M) estimator [49] and shown in Fig. 6.b. A multivariate log-rank test [50] is conducted to compare their survival outcomes, revealing a statistically significant difference (p = 0.00056) among the groups. Hazard ratios for these patient subgroups are calculated using the Cox proportional regression model [51], and are presented in Fig. 6.c. The results reveal that S1 exhibits statistically significantly worse survival compared to other patients (hazard ratio (hr) = 2.43, 95% confidence interval (CI): 1.43-4.11, p = 0.0009). S2 also demonstrates significantly worse survival (hr = 1.67, 95% CI: 1.06-2.65, p = 0.03). Conversely, S7 exhibits statistically significantly better survival (hr = 0.51, 95% CI: 0.32-0.82, p = 0.005).

#### 2.4.2 BiGraph provides complementary information to standard clinical characteristics

It is natural to ask how the population graph – constructed based on the similarities between patients’ TME patterns – relates to clinical characteristics. Does it echo the information already encoded in clinical variables, or does it provide new information and insights? To answer this question, we color the nodes of the population graph with clinical subtypes (Fig. 7.a), tumor stage (Fig. 7.b), tumor grade (Fig. 7.c), and patients’ age (Fig. 7.d). The figures demonstrate that close nodes can have different colors, implying that patients with similar TMEs do not necessarily have the same clinical attributes. Moreover, we calculate the average inter-patient similarity for each group of patients stratified based on clinical variables or based on BiGraph (see Table 1). The results confirm this observation, as the average inter-patient similarity within the same clinical group is similar to (not higher than) that of the entire heterogeneous set. Yet, there is an exception: Her2+ patients show a higher average interpatient similarity (0.43) compared to the entire cohort (0.18). Spearman’s correlation scores [52] are used to assess the alignment between BiGraph-derived subgroups and clinical groups, revealing that they are generally not aligned with each other well (see Fig. 7.e). Yet, certain correlations do exist. Specifically, S1 is strongly positively correlated with HER2+ patients, and negatively correlated with HR+/HER2-. The group S7, on the other hand, exhibits positive correlations with HR+/HER2-, grade 1, and grade 3, while being negatively correlated with grade 3. Additionally, S2 and S4 are positively correlated with TNBC.

**Fig. 7:**
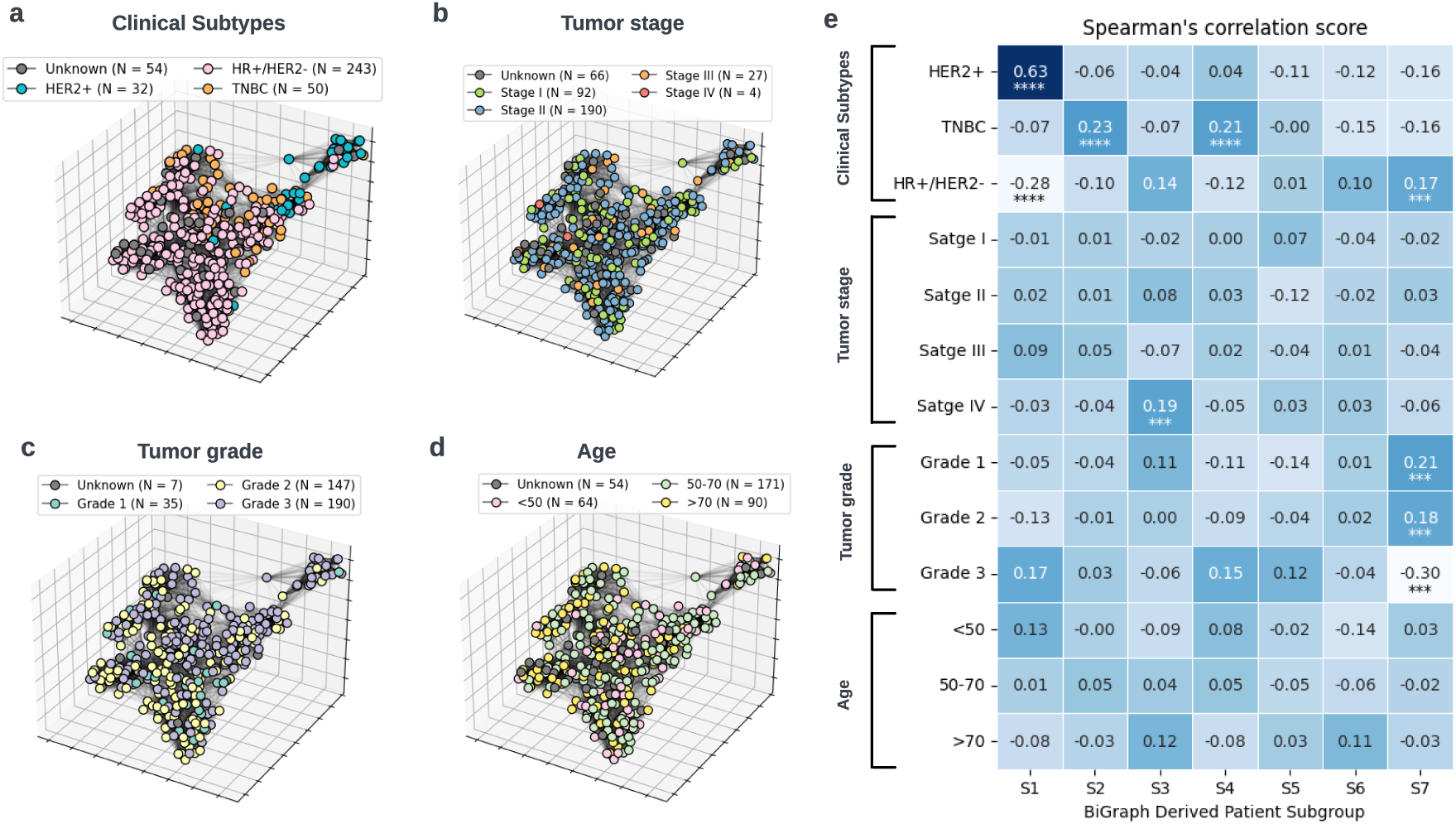
Correlation between clinical characteristics and BiGraph-derived subtypes. **(a)** Population Graph with nodes painted by clinical subtypes. **(b)** Population Graph with nodes painted by tumor stage. **(c)** Population Graph with nodes painted by tumor grade. **(d)** Population Graph with nodes painted by age group. **(e)** Spearman’s correlation scores between clinical characteristics and BiGraph-derived subgroups. ∗ ∗ ∗∗: p-value *<* 1e-4; ∗ ∗ ∗: p-value *<* 1e-3.

**Table 1:**
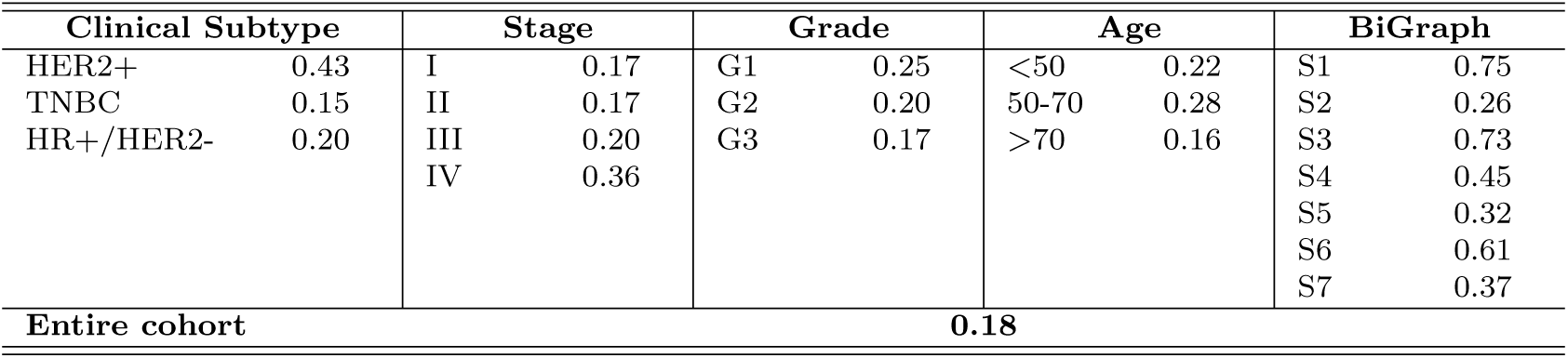
Average inter-patient similarities of various clinical groups and BiGraph-derived subgroups.

A crucial observation is that the intersection sets between BiGraph-derived subtypes and clinical subtypes reveal unique survival patterns in some cases. For instance, HER2+ patients (N = 32) exhibit worse survival compared to others in the discovery set. Further stratification using S1 identifies a subset (N = 19) with even worse survival, and a complement subset (N = 13) in the HER2+ patients with slightly better survival (Fig. 8.a). Similarly, S2 divides TNBC patients (N = 50) into a subset (N = 16) with exceptionally worse survival and a complement set (N = 34) (Fig. 8.b). Furthermore, HR+/HER2-patients (N = 243) display better survival compared to others, while its intersection with S7 patients (N = 76) shows even better survival (Fig. 8.c). These results suggest that the BiGraph-derived subtyping system provides complementary information for the clinical subtyping system, enhancing its risk stratification capacity beyond that achieved by the clinical subtyping system alone.

**Fig. 8:**
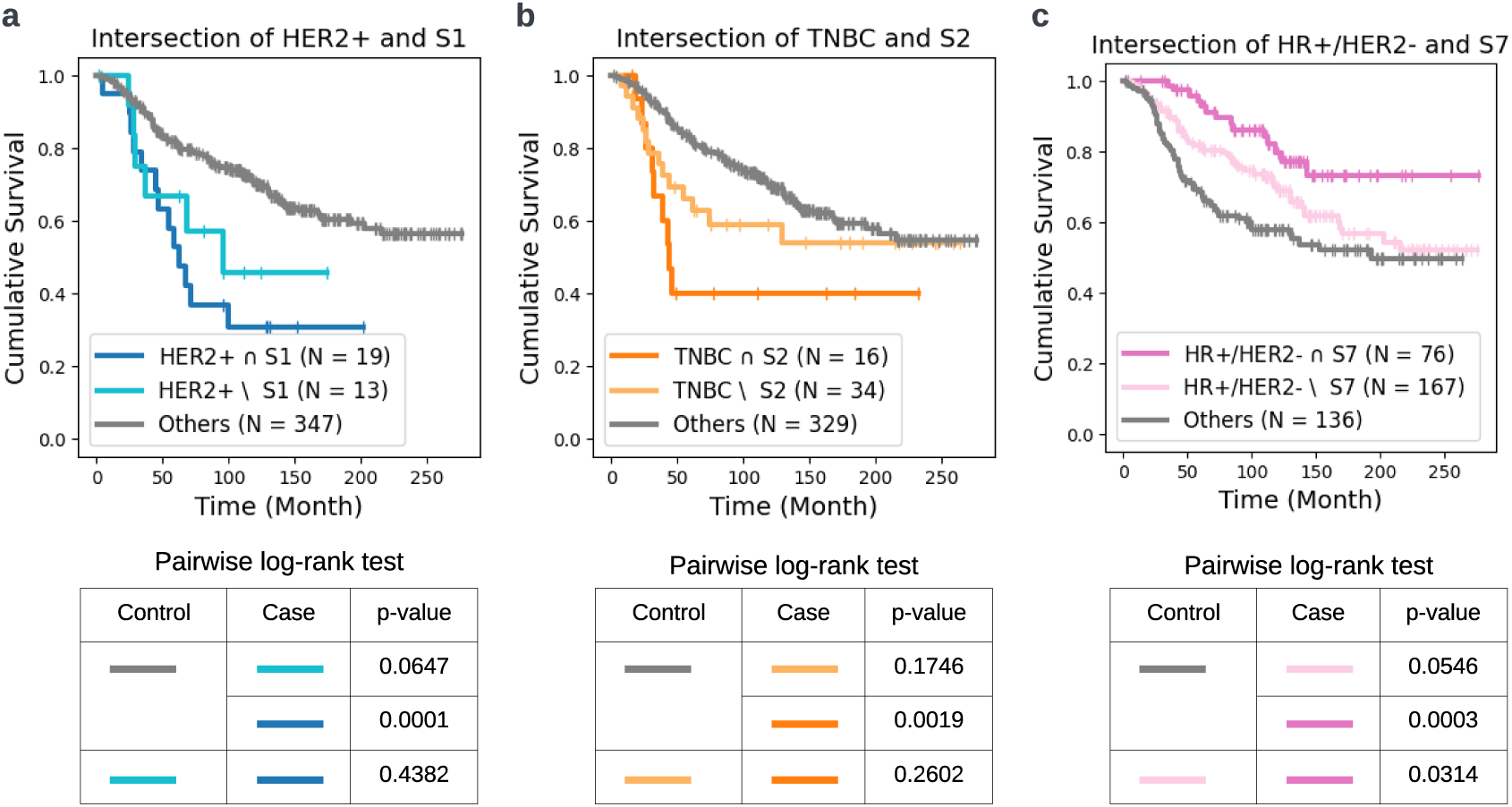
BiGraph enhances the risk stratification of clinical subtype. **(a)** K-M survival plots of the intersection of HER2+ and S1 patients; the complement of S1 in HER2+ patients; and all other patients. **(b)** K-M survival plots of the intersection of TNBC and S2 patients; the complement of S2 in TNBC patients; and all other patients. **(c)** K-M survival plots of the intersection of HR+/HER2- and S7 patients; the complement of S7 in HR+/HER2- patients; and all other patients.The pairwise log-rank tests are used to compare the survivals of three groups of patients in each case.

### 2.5 BiGraph unveils prognosis-relevant TME patterns

An important feature of BiGraph – distinguishing it from alternative, more opaque data-driven methods – is its explainability. The construction of the population graph relies on comparing the histogram of TME patterns in patients via the soft WL subtree kernel, which allows us to directly identify the most characteristic patterns for each patient subgroup. Furthermore, the prognostic impacts of these characteristic patterns can be systematically investigated.

#### 2.5.1 Patient subgroups are characterized by unique TME patterns

To identify characteristic patterns of different survival profiles, the expression of TME patterns within and outside a patient subgroup are compared using the Hodges–Lehmann statistic [53]. A TME pattern is deemed “characteristic” in a patient subgroup if its Hodges–Lehmann statistic surpasses 50% of the maximum value (see Section 5.5, Extended Data Fig. 4). Results indicate that distinct patient subgroups exhibit characteristic patterns with diverse cellular compositions and organizations (see Fig. 9). Representative examples of these characteristic patterns are exhibited in Fig. 10. Although subtrees show the hierarchical structure of cells in an abstract manner, they do not intuitively convey the spatial cellular organization. For interpretation purposes, a corresponding cellular neighborhood centered at the subtree root is displayed in Fig. 10, where the neighborhood boundary is determined by cells’ contributions to the subtree feature embedding (see Section 5.3.1, Equation (3)).

**Fig. 9:**
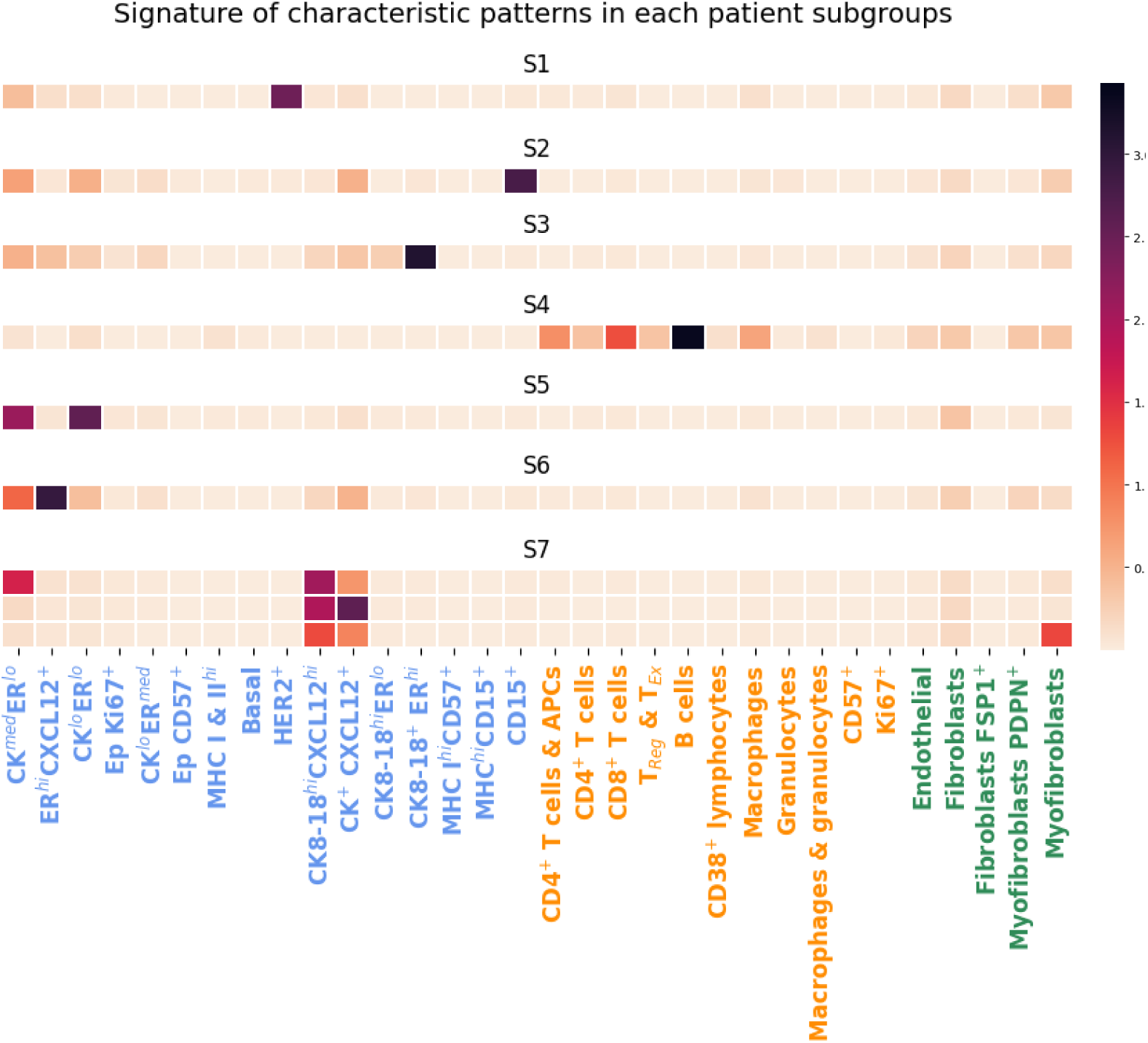
Signature of characteristic patterns in each patient subgroup. Each row shows the signature (i.e., averaged feature embedding) of a characteristic pattern. The x-axis indicates cell phenotype and x-axis text color indicates the cell type category: blue: tumor cells; orange: immune cells; green: stromal cells.

**Fig. 10:**
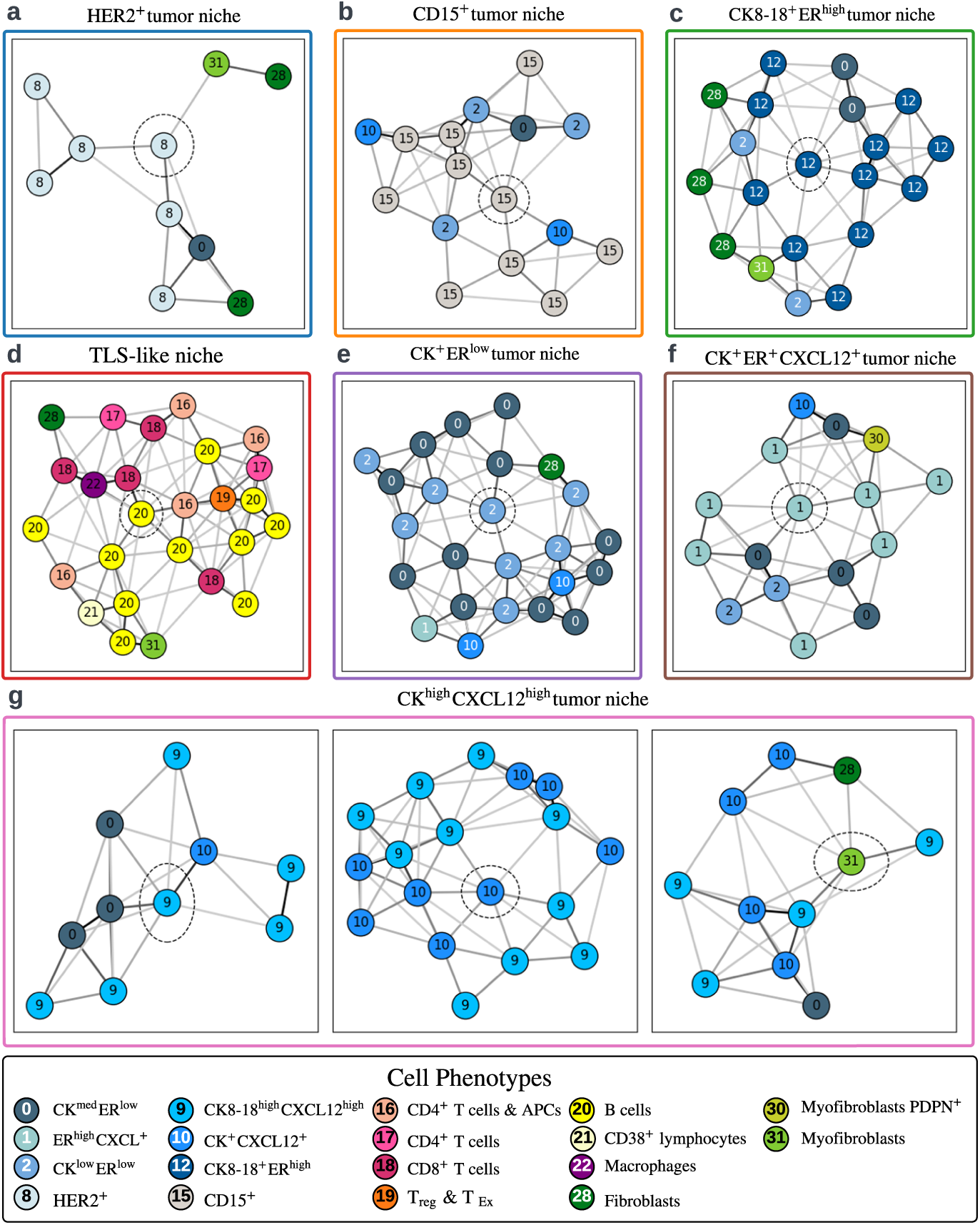
Representative examples of characteristic patterns in each patient subgroup. To intuitively show the spatial organization of cells, the corresponding cellular neighborhoods of subtrees are shown, where dashed circles indicate the subtree root. **(a)** HER2^+^ tumor niche, characteristic in S1; **(b)** CD15^+^ tumor niche, characteristic in S2; **(c)** CK8-18^+^ER^high^ tumor niche, characteristic in S3; **(d)**: TLS-like niche, characteristic in S4; **(e)** CK^+^ER^low^ tumor niche, characteristic in S5; **(f)** CK^+^ER^+^CXCL12^+^ tumor niche, characteristic in S6; **(g)** CK^high^CXCL12^high^ tumor niche, characteristic in S7.

Patient subgroup S1, exhibiting the worst survival outcome, is distinguished by “HER2^+^ tumor niche” patterns, showing enrichment of HER2^+^ tumor cells surrounded by fibroblasts and myofibroblasts. In S2, a patient subgroup exhibiting a statistically significantly worse survival outcome, “CD15^+^ tumor niche” patterns are most characteristic, featuring enriched CD15^+^ tumor cells intermixed with some other tumor cells. S3, a relatively small and homogeneous patient subgroup, is characterized by an “enriched CK8-18^+^ER^high^ tumor niche”. The most distinctive pattern within S4 closely resembles the well-known tertiary lymphoid structure (TLS) [54], and is termed a “TLS-like niche”. This pattern encompasses a variety of immune cells, including B cells (predominantly represented), CD4^+^ T cells, CD8^+^ T cells, regulatory T cells (Treg), macrophages, and CD38^+^ lymphocytes. The most characteristic pattern in S5 is a “CK^+^ER^low^ tumor niche”, exhibiting a mixture of CK^med^ER^low^ and CK^low^ER^low^ tumor cells. S6 is characterized by a “CK^+^ER^+^CXCL12^+^ tumor niche” pattern, featuring the co-localization of CK^med^ER^low^ and ER^high^CXCL12^+^ tumor cells. S7, the patient subgroup exhibiting the best survival outcome, has technically three characteristic patterns, while they exhibit a common feature: a mixture of CK8-18^high^CXCL12^high^ cells and CK^+^CXCL12^+^ cells. Therefore, these three patterns are consolidated into one, named a “CK^high^CXCL12^high^ tumor niche”.

#### 2.5.2 Prognostic values of the characteristic TME patterns

The results above showcase the most characteristic TME patterns in each subgroup, but these do not necessarily imply associations with prognosis. This naturally leads to the exploration of potential associations between these characteristic TME patterns and patient survival outcomes, which we explore next. The prognostic impacts of a specific TME pattern are assessed by categorizing patients into “positive” and “negative” groups based on the proportion of that pattern in a patient. A 1% threshold is employed for patient stratification, the same as the cut-off point used in the definition of hormone receptor (HR) status [55]. For each characteristic pattern, survival outcomes of positive and negative patients are compared using the log-rank test. Results indicate that the HER2^+^ tumor niche, characteristic of S1, is associated with worse survival (p = 0.00031) (Fig. 11.a). In contrast, the CK^high^CXCL12^high^ tumor niche, characteristic of S7, is associated with better survival (p = 0.025) (Fig. 11.g). For patterns characteristic in S5 and S6, positive patients exhibit slightly better survival outcomes than negative patients, although the differences are not statistically significant. These results are broadly consistent with the risk stratification provided by the seven patient subgroups – S1 patients display significantly worse survival outcomes, and the most characteristic TME pattern proves to be a negative prognostic factor. Conversely, S7 patients exhibit significantly better survival outcomes, with the most characteristic TME pattern emerging as a positive prognostic factor.

**Fig. 11:**
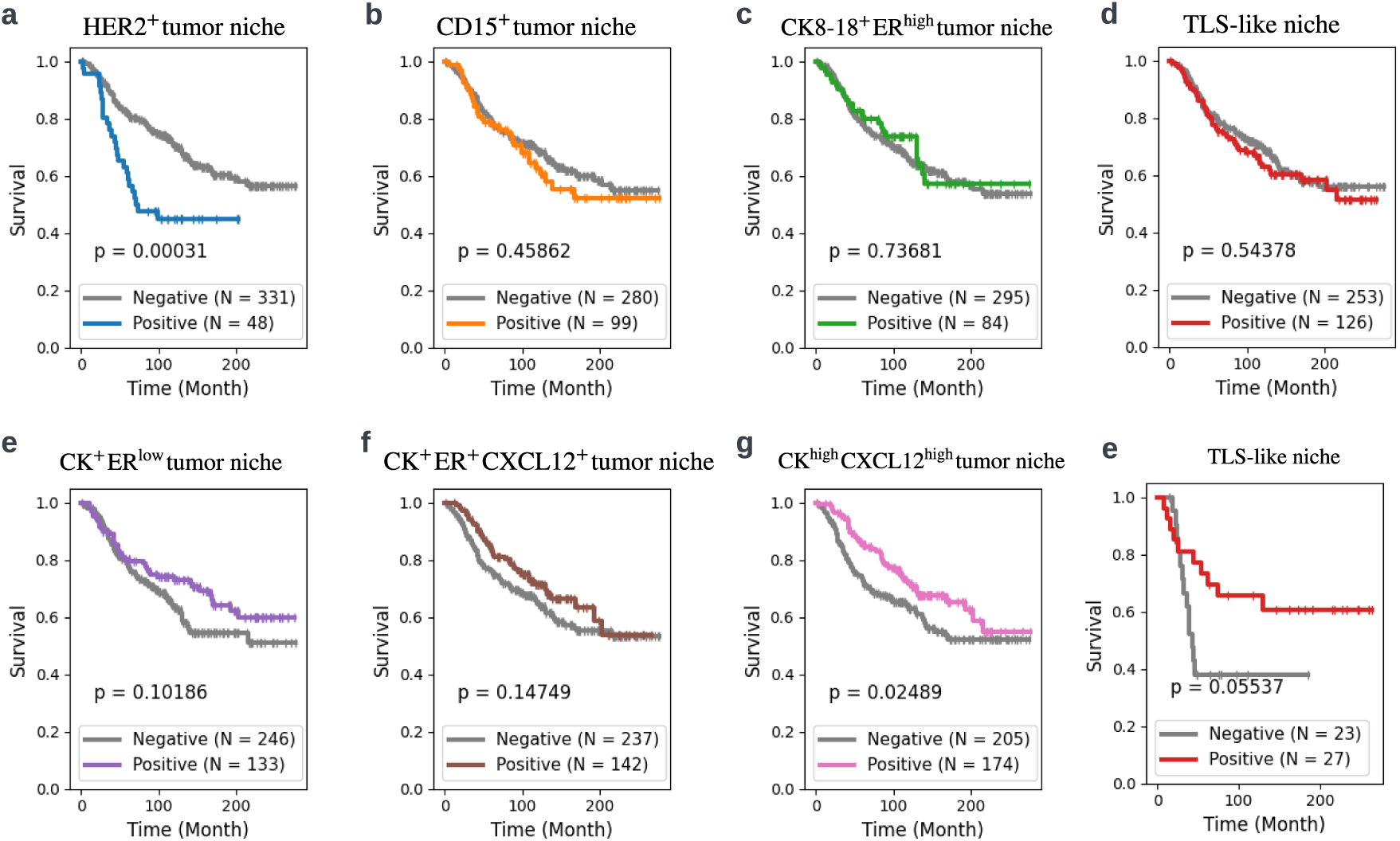
Prognostic impacts of the characteristic TME patterns. Patients are categorized into “positive” and “negative” groups based on the proportion of a designated TME pattern with 1% as the threshold. Their survivals are compared by log-rank test, and the resulting p-values are indicated in each corresponding figures.**(a)** risk stratification of HER2^+^ tumor niche; **(b)** risk stratification of CD15^+^ tumor niche; **(b)** risk stratification of CK8-18^+^ER^high^ tumor niche; **(d)**risk stratification of TLS-like niche; **(e)** risk stratification of CK^+^ER^low^ tumor niche; **(f)** risk stratification of CK^+^ER^+^CXCL12^+^ tumor niche; **(g)** risk stratification of CK^high^CXCL12^high^ tumor niche; **(e)** risk stratification of TLS-niche, specifically in TNBC patients.

Moreover, since TNBC is a especially aggressive and challenging breast cancer subtype, we further examine the prognostic values of these patterns specifically within TNBC patients. Interestingly, the TLS-like niche, characterized by densely aggregated B cells surrounded by CD4^+^ T cells, CD8^+^ T cells, macrophages, and stromal cells, shows no prognostic value across the entire discovery set (N = 379) with heterogeneous clinical subtypes (Fig. 11.d). Yet, in the TNBC subset (N = 50), patients with TLS-like niches present in their TMEs do exhibit better survival than negative patients (Fig. 11.e). Although this difference is not statistically significant, likely due to the small cohort size, the large divergence of the two survival curves is noteworthy.

To better understand the prognostic impacts of these prognosis-relevant TME patterns, we investigate their correlation with standard clinical variables, such as tumor grade, stage, metastasis, and clinical subtypes. Results suggest that the expression of HER2^+^ tumor niche per patient is significantly higher in grade 3 patients (see Fig. 12.a) and in Her2+ patients (see Fig. 12.d). In contrast, the expression of CK^high^CXCL12^high^ tumor niche is significantly lower in grade 3 patients (see Fig. 12.e) but significantly higher in HR+/HER2-patients (see Fig. 12.f). These findings highlight the association between HER^+^ tumor niches with more aggressive clinical features (i.e., higher grade and HER2+ subtype), and the association between CK^high^CXCL12^high^ tumor niches and less aggressive features (i.e. lower grade and HR+/HER2-subtype). For TLS-like niches, the analysis is constrained to TNBC patients as it only shows links to prognosis in this subset. Results show no significant correlation between TLS-like niches and grade, stage, or metastasis (see Fig. 12.j-e).

**Fig. 12:**
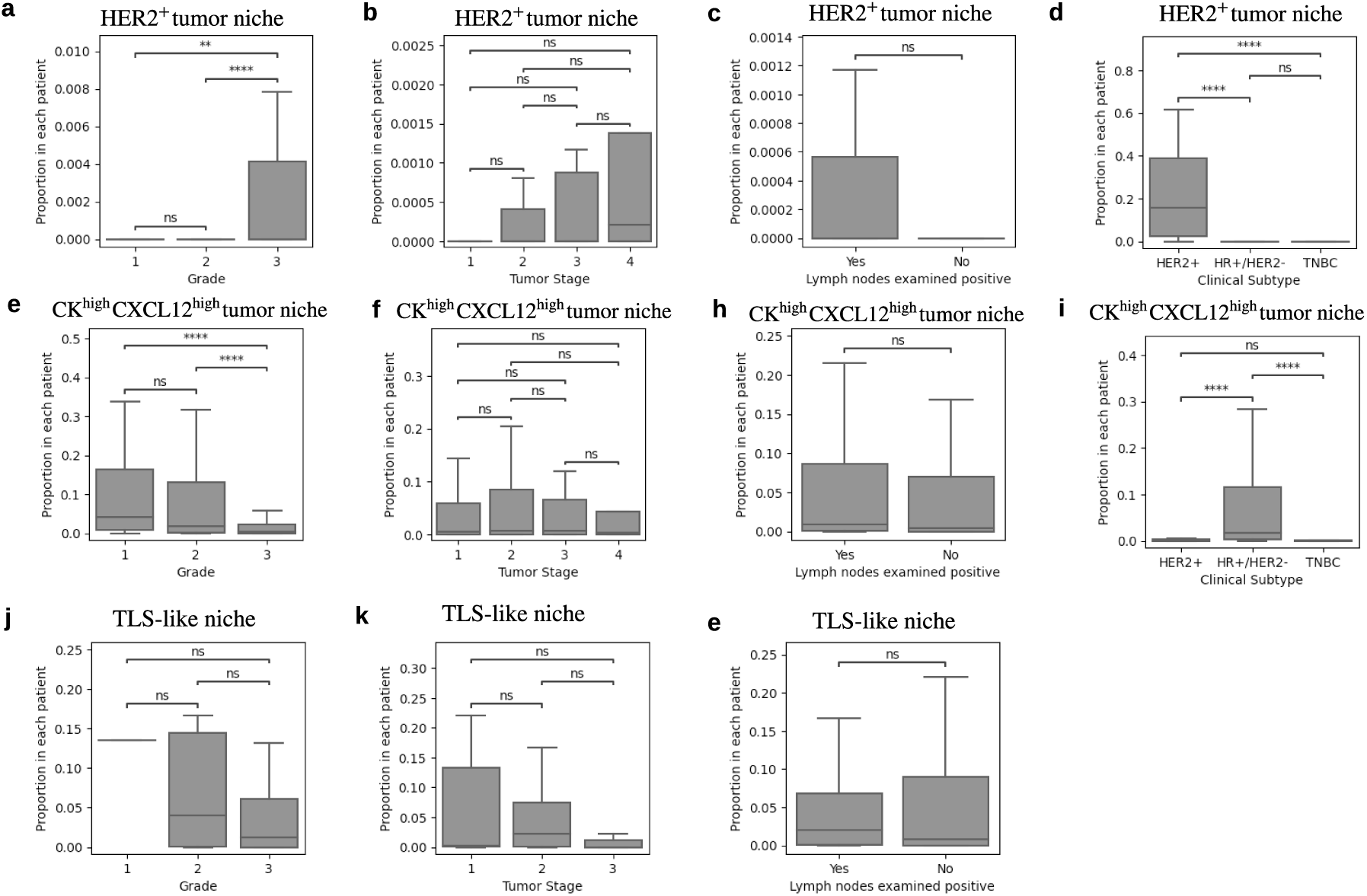
Correlation between prognosis-relevant TME patterns and clinical variables. **(a) - (d)** The boxplots of the proportion of HER2^+^ tumor niche in different tumor grades, tumor stages, metastasis status, and clinical subtypes. **(e) - (f)** The boxplots of the proportion of CK^high^CXCL12^high^ tumor niche in different tumor grades, tumor stages, metastasis status, and clinical subtypes. **(j) - (e)** The boxplots of the proportion of TLS-like niches in different tumor grades, tumor stages, and metastasis status. For TLS-like niches, only TNBC patients are included. Outliers are not shown in the boxplots. Mann–Whitney U test is used to compare proportions. ∗ ∗ ∗∗: p-value *<* 1e-4; ∗ ∗ ∗: p-value *<* 1e-3. ∗∗: p-value *<* 1e-2.

### 2.6 Validation and generalization

As detailed in Section 2.1, a subset comprising 200 patients is reserved from the 579 patients curated by Danenberg et al. [16], denoted as the “inner validation set”. Additionally, an external breast cancer patient cohort curated by Jackson et al. [17], referred to as the “external validation set-1”, and an additional external TNBC patient cohort curated by Wang et al. [47], referred to as the “external validation set-2”, are utilized to evaluate BiGraph’s cross-study generalization capacity. Importantly, these validation sets allow us to explore generalization under different settings: the inner validation set provides generalization results to data collected under the same protocol as those employed in the discovery set, whereas the other two external validation sets involve data collected independently: the external validation set employs a distinct antigen panel and phenotype system compared to the discovery set. Moreover, external validation set-2 is a clinical trial, allowing us to validate the predictive value of specific TME patterns. This external validation setting adds complexity to the validation process, while on the other hand provides a stronger demonstration of the applicability of BiGraph in real-world scenarios – as standardized antigen panels are not yet common in practice. To address this challenge, we employ a mapping approach to align the cell phenotyping in the external validation set with that of the discovery set. This alignment is achieved by identifying shared antigens between both sets, as indicated by shaded regions in Fig. 3.f. To be more precise, every cell in the external validation set is mapped to the closest cell phenotype cluster in the discovery set, where the cell-to-cell phenotypic distance is given by the Euclidean distance of their expressions of the shared antigens (see Section 5.7.1, and the cell phenotyping alignment quality is shown in Extended Data Fig. 5). With the cell phenotyping system of the external validation set aligned, we validate the major results found in the discovery set in both inner- and external validation sets.

#### 2.6.1 Validation of the patient risk stratification

As detailed in Section 2.4.1, the community detection algorithm identifies seven patient subgroups from the population graph of the discovery set with significantly distinct survivals. In the validation stage, every unseen new patient (with their corresponding cellular graph) is mapped to one of the seven patient subgroups using a *k*-nearest neighbor (*k*-NN) rule [56]. The similarity between a new patient and patients in the discovery set is measured by the Soft-WL subtree kernel function defined on the discovery set (see Section 5.7.3). In this way, seven mapped patient subgroups can be identified in the inner- and external-validation sets respectively, denoted as S1′, S2′, etc., where these “prime” counterparts denote “validation”.

We analyze the survival outcomes of the seven mapped patient subgroups in both inner validation (Fig. 13.a-c) and external validation set-1 (Fig. 13.d-f). Note that validation of risk stratification on external validation set-2 is not applicable, since the long-term follow-up data is unavailable. The multivariate log-rank test is used to compare K-M survival plots of the seven patient subgroups. The survival outcomes of the seven mapped patient subgroups do not have a statistically significant difference (p = 0.24, Fig. 13.b) in the inner-validation set. On the other hand, there is a statistically significant distinction among their survival outcomes in the external validation set (p = 0.007, Fig. 13.e). By comparing the hazard ratios of the seven mapped subgroups with the subgroups found in the discovery set (Fig. 6.c), we observe a consistent trend from S1 to S7, which provides support for the reproducibility of our findings. Furthermore, our validation results confirm that S1^′^ in the external validation set exhibits significantly worse survival (p*<*0.05), consistent with the prognosis of S1 in the discovery set. Similarly, S7^′^ in the external validation set demonstrates better survival outcomes (p*<*0.05), mirroring the survival pattern of S7 observed in the discovery set. In the inner validation set, S1′ has a comparatively high hazard ratio, and S7′ has a comparatively low hazard ratio, while both lack statistical significance.

**Fig. 13:**
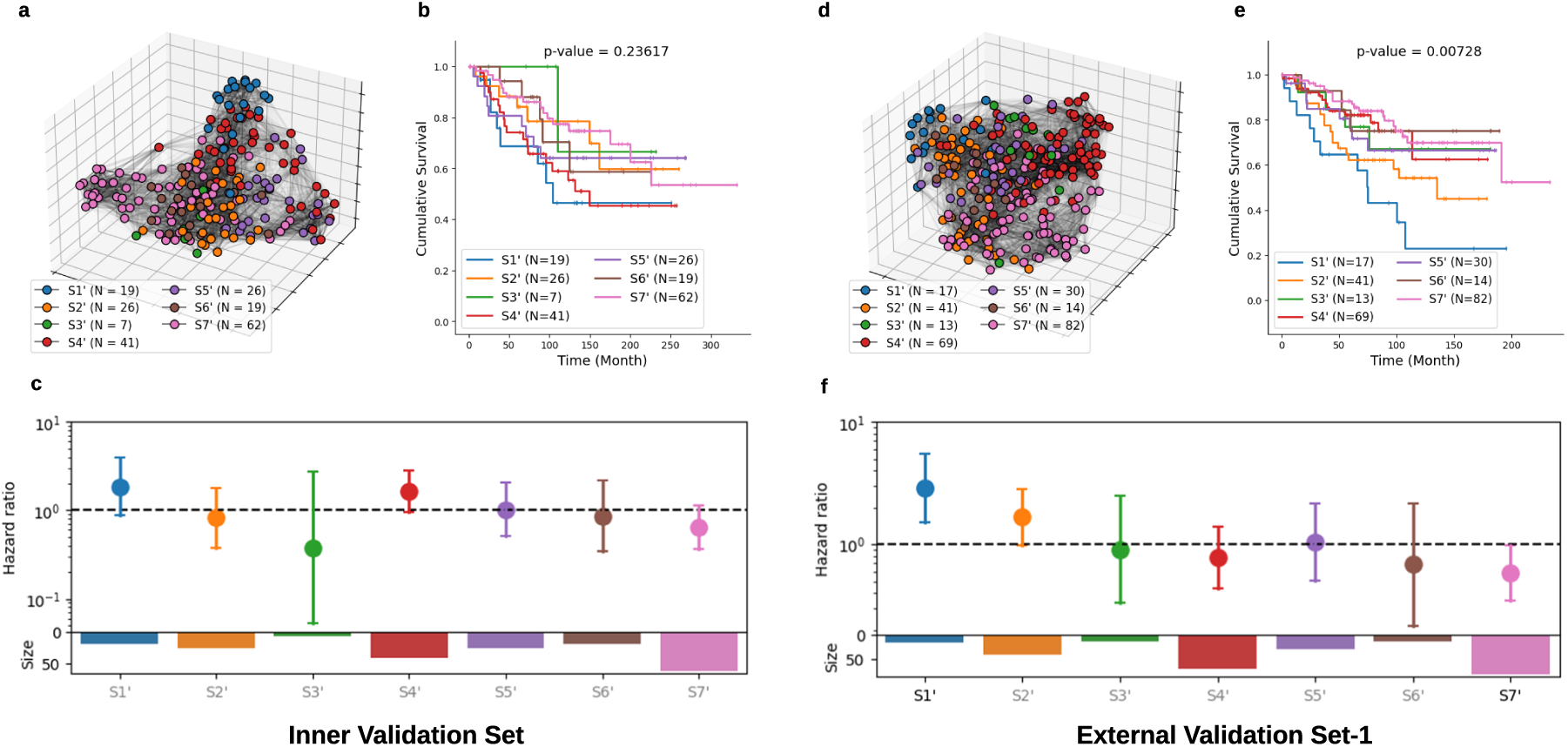
Validation of the risk stratification results. **(I)** Results in inner validation set (N=200). **(E)** Results in external validation set (N=266). **(a)** Population graph with nodes painted by estimated patient subgroup labels; **(b)** K-M estimation of the survival functions of seven mapped patient sub-groups. A multivariate log-rank test is used to compare them, and the p-value is indicated in the title.; **(c)** Hazard ratios of the seven mapped patient subgroups. The text color (grey vs. black) distinguishes between statistically significant and non-significant associations (black: statistically significant). The size (i.e., the number of patients) of each subgroup is presented in the corresponding barplot.

Moreover, and importantly, we find that the intersection set between a clinical subtype and a BiGraph-derived subgroup exhibits exceptionally better (or worse) survivals in some cases (Fig. 7). We validate these results on the two validation sets. While the intersection of HER2+ patients and S1^′^ patients (Fig. 14.a and d), and the intersection of TNBC patients and S2^′^ patients (Fig. 14.b and e), show contradictory results in the validation sets, the intersection between HR+/HER2-patients and S7 patients (Fig. 14.c and f) indeed has an exceptionally better survival compared with the complement of S7 in HR+/HER2- patients and patients with other clinical subtypes, though this is not statistically significant. The lack of significance might be due to several factors, including the variant nature of the patient cohort, differences in data acquisition protocols (e.g., different antigen panels), potential distribution shifts, and the smaller size of the validation sets, all of which pose substantial changes to validating findings from the discovery set.

**Fig. 14:**
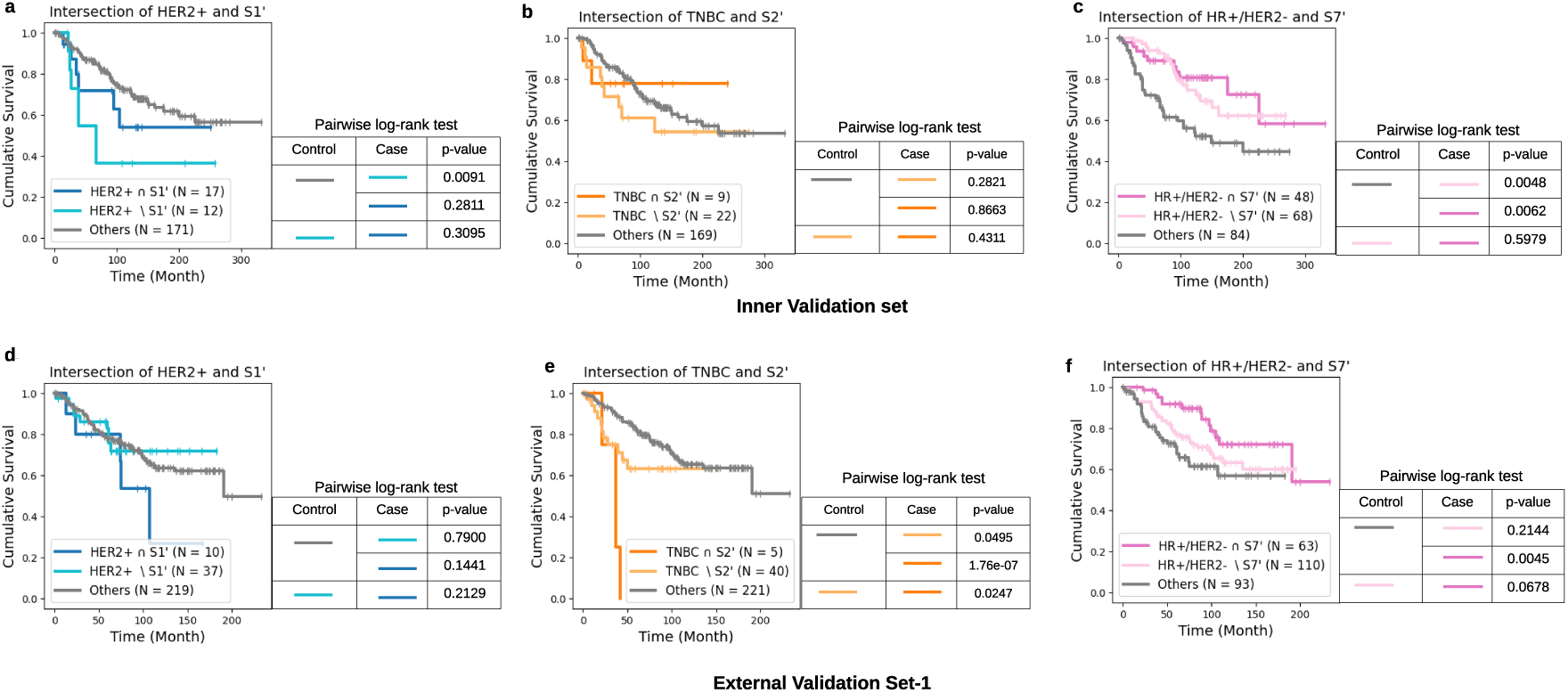
Validation of the enhancement of risk stratification that BiGraph provides to clinical subtyping. **(a)** and **(d)** Survival plots of the intersection of HER2+ and S1^′^ patients; the complement of S1^′^ in HER2+ patients in inner validation set, and external validation-1; and all other patients. **(b)** and **(e)** Survival plots of the intersection of TNBC and S2^′^ patients; the complement of S2^′^ in TNBC patients; and all other patients in inner validation set, and external validation-1. **(c)** and **(f)** Survival plots of the intersection of HR+/HER2- and S7^′^ patients; the complement of S7^′^ in HR+/HER2- patients; and all other patients in the inner validation set, and external validation-1.The pairwise log-rank tests are used to compare the survivals of three groups of patients.

#### 2.6.2 Validation of the prognostic values of TME patterns

As detailed in Section 2.5, we find two prognostic-relevant TME patterns in the discovery set: the HER2^+^ tumor niche (associated with worse survival) and the CK^high^CXCL12^high^ tumor niche (associated with better survival). Moreover, while the TLS-like niche is not associated with distinct prognosis across the entire discovery set, it is linked to better survival in TNBC patients. We validate the prognostic impacts of HER2^+^ tumor niche and CK^high^CXCL12^high^ tumor niche in both inner validation set and external validation set-1. The prognostic value of the TLS-like niche is validated in TNBC patients of the inner validation set, external validation set-1, and external validation set-2 (which only contains TNBC patients).

To estimate the expression of the two TME patterns of interest in unseen patients, we map the subtrees derived from their cellular graphs to the closest TME pattern (see Section 5.7.2). The signatures of each TME pattern in the discovery set and validation sets are qualitatively compared to assess the TME pattern mapping quality (see Extended Data Fig. 6). Furthermore, the distributions of each TME pattern’s proportion across discovery and validation sets are compared to identify any potential distribution shifts (see Extended Data Fig. 7). Patients in the validation sets are categorized as “positive” or “negative” based on the expression of the studied TME pattern, with a 1 % proportion as the cut-off point as before. Notably, while the discovery set and external validation set-1 consist of observational series where patients received standard care, external validation-2 is a clinical trial where patients received chemotherapy with or without immune therapy (C vs. C&I). Since long-term follow-up data is unavailable, pathological response post-treatment serves as a surrogate for prognosis. In this case, we compare the proportion of TLS-like niches for patients who achieved pathologically complete response (pCR) versus those with residual disease (RD).

Results show that HER2^+^ tumor niche positive expressed patients indeed exhibit worse survival outcomes than negatives, in both the inner validation set (p = 0.0002) and external validation set (p = 0.097), with the distinction being statistically significant only in the inner validation set (Fig. 15.a and d). On the other hand, CK^high^CXCL12^high^ tumor niche-positive patients demonstrate better survival outcomes than the remaining in both inner validation set (p = 0.055) and external validation set (p = 0.007), with the distinction being statistically significant only in the external validation set ((Fig. 15.b and e). For the TLS-like niche in TNBC patients, the inner-validation set and external validation set-1 show a slight trend towards better survivals for TLS-like niche-positive patients, though the differences are not statistically significant, likely due to the small size of 31 and 45 TNBC patients in these sets (Fig. 15.c and f). In external validation set-2, the proportion of TLS-like niche is assessed in the tissue samples collected before (i.e., pre-treatment samples) and at the early stage of the treatment (i.e., on-treatment samples) for patients receiving chemotherapy alone (C) and chemotherapy combined with immune therapy (C&I). The results indicate that in both pre-treatment and on-treatment samples the proportion of TLS-like niche is statistically significantly higher in patients with pCR (better outcome) compared to those with RD (worse outcome) (Fig. 15.g and h). Specifically, a higher presence of TLS-like niches in on-treatment samples strongly indicated better outcomes for patients receiving C&I (Fig. 15.j).

**Fig. 15:**
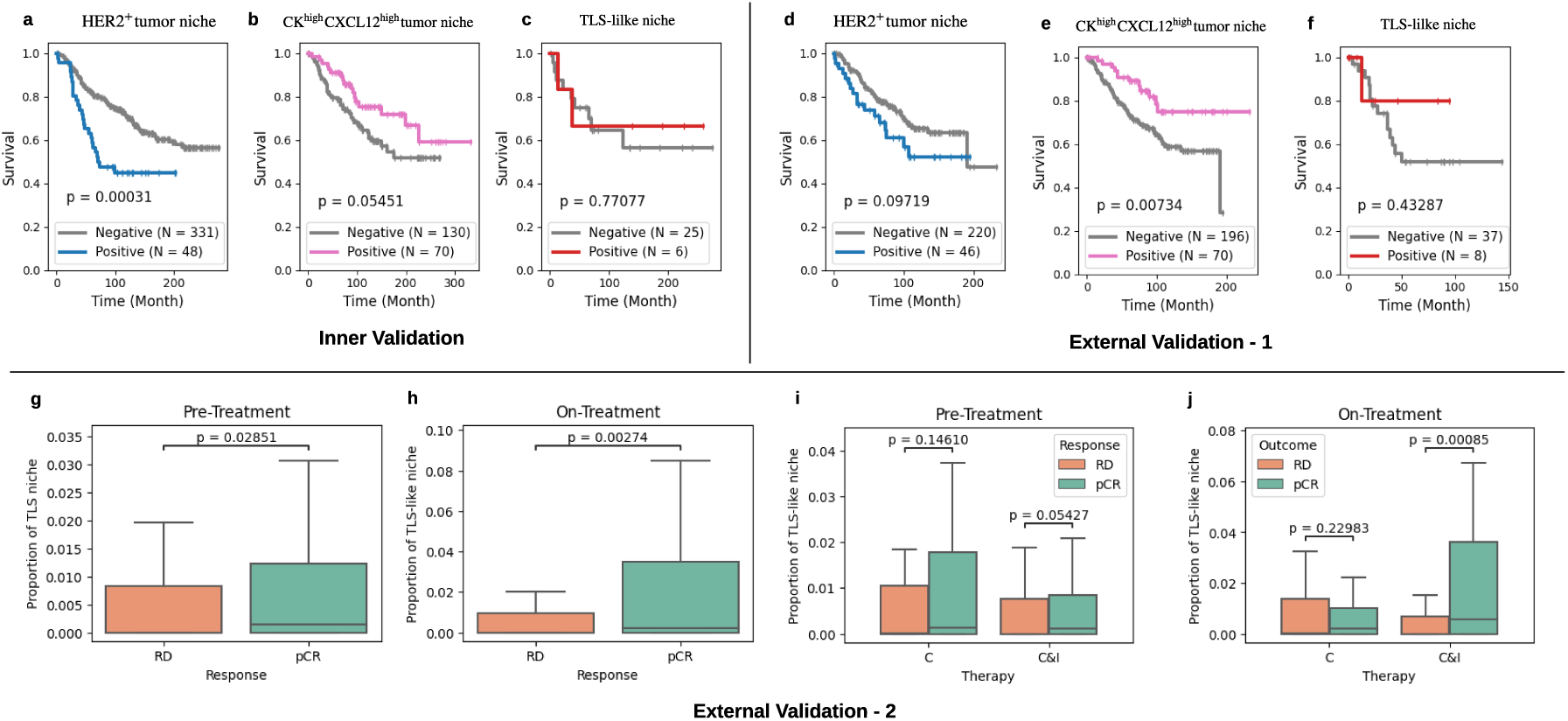
Validation of the prognostic impacts of TME patterns. **(a)** K-M survival plots of patients with (proportion ≥ 1%) and without HER2 tumor niche in inner-validation set. **(b)** K-M survival plots of patients with and without CK^high^CXCL12^high^ tumor in inner-validation set. **(c)** K-M survival plots of patients with and without TLS-like niche in TNBC patients of the inner-validation set. **(d) - (f)** Same analysis as **(a) - (c)**, but on external validation-1. **(g-h)** The proportion of TLS-like niches in the pre- and on-treatment samples for patients with residual disease (RD) and pathologically complete response (pCR) in external validation set-2. (i-j) Boxplot of the proportions of TLS-like niches in pre- and on-treatment samples for patients with RD or pCR in external validation set-2, stratified by treatment type (chemotherapy (C) vs. chemotherapy and immunotherapy (C&I)). Mann–Whitney U test is used to test for differences in samples. Outliers are not shown.

Additionally, the relationship between these two prognosis-relevant TME patterns and tumor grade is validated. HER2^+^ tumor niche is statistically significantly over-expressed in grade 3 patients in the inner validation set (Fig. 16.a), consistent with the results on the discovery set. However, there are no significant differences in the external validation set (Fig. 16.c). On the other hand, both inner validation set and external validation set results confirm that CK^high^CXCL12^high^ tumor niche is statistically significantly under-expressed in grade 3 patients (Fig. 15.b and d), supporting our findings.

**Fig. 16:**
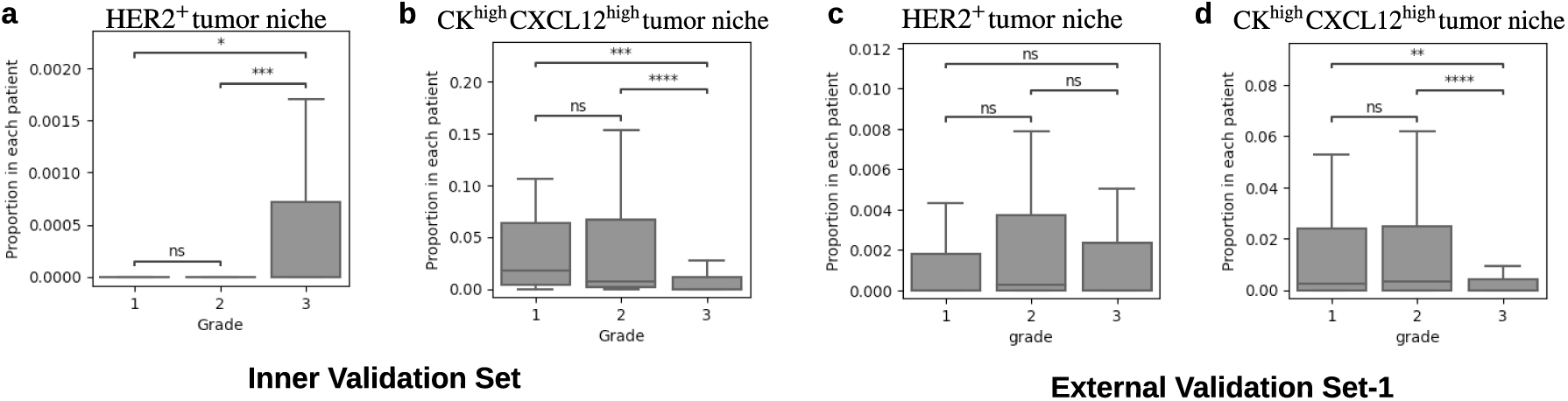
Validation of the correlation between TME patterns and tumor grade. **(a) and (c)** Boxplot of the proportion of HER2^+^ tumor niche per patient for tumor grade 1, 2, and 3 in the inner validation set and external validation set-1; **(b)** and **(d)** Boxplot of the proportion of CK^high^CXCL12^high^ tumor niche per patient for tumor grade 1, 2, and 3 in inner validation set in inner validation set and external validation set-1. Outliers are not shown in the boxplots. Mann–Whitney U test is used to compare proportions. ∗ ∗ ∗∗: p-value *<* 1e-4; ∗ ∗ ∗: p-value *<* 1e-3. ∗∗: p-value *<* 1e-2.

### 2.7 Comparison with alternative methods

The primary insight behind BiGraph is to establish a link between patient-specific cellular features (based on the spatial distribution of cell types) and the population-level distribution by utilizing a graph kernel function. In the previous discussions, we introduced the Soft-WL subtree kernel, which measures inter-patient similarity by assessing the abundance of TME patterns representing local structures within TMEs. This naturally prompts the question: are there alternative approaches to measuring inter-patient similarity giving similar or novel discoveries? We keep the process of construction of a population graph, community detection, and survival analysis consistent with previous sections, but use alternative methods to calculate inter-patient similarities.

1. **Abundance of cell types:** In this approach, each patient is characterized by a histogram of cells, and inter-patient similarity is determined by the normalized inner product of corresponding histograms. This approach disregards spatial information and inter-cellular interactions. The community detection algorithm identifies seven patient subgroups in the population graph, denoted as S1 to S7. Notably, only S1 exhibits significantly worse survival (see Fig. 17.a). The S1 subgroup identified here and that found by the Soft-WL subtree kernel reveals a large overlap, with an intersection over union (IoU) score of 0.85. Furthermore, S1 identified here is characterized by an over-presentation of HER2^+^ cells, aligning with the HER2^+^ tumor niche, characteristic in S1 found by Soft-WL subtree kernel. However, the S7 subgroup identified by the Soft-WL subtree kernel, characterized by TME patterns involving multiple types of cells, does not align with any of the patient subgroups identified using the abundance of cell types.
2. **Pairwise proximity of cell types:** In this approach, each patient is characterized by the average proximity score – which increases as the spatial distance decreases – between any possible pairs of cell phenotypes (see Equation (6)). While this method considers spatial information to some extent, it is limited to pairwise relationships. The resulting population graph does not demonstrate any clear clusters, and the community detection algorithm identifies six patient subgroups, none of which exhibit distinct survival outcomes (see Fig. 17.b).
3. **Abundance of TME categories:** As described in Section 2.2, the 66 TME patterns are grouped into four main categories: tumor niches, immune niches, stromal niches, and interface niches based on the predominant cell phenotypes within each subtree. In this case, each patient is characterized by a histogram of these four TME categories, and inter-patient similarity is determined by the cosine similarity of corresponding histograms. This approach is a coarser version of the BiGraph method, presented in the previous sections, since fine-grained differences among TME patterns are omitted. The resulting population graph unveils six patient subgroups, but none of these exhibit distinct survival outcomes (see Fig. 17.c).
4. **WL subtree kernel:** The Soft-WL subtree kernel, presented in this work, measures inter-patient similarity by identifying identical TME patterns across patients, each corresponding to a cluster of similar subtrees. In contrast, the classic WL subtree kernel [43] iteratively identifies *isomorphic* subtrees across graphs (see Section 5.8). We explore two formulations of the WL subtree kernel: one that considers isomorphic subtrees found in all previous iterations and accumulates the similarity (see Equation (8)), and another that only considers isomorphic subtrees found at the last iteration (see Equation (9)). Results indicate that the former formulation yields almost the same results as characterizing patients by the abundance of cell types (see Extended Data Fig. 8). This is attributed to the stringent criteria of isomorphic comparison. As iteration progresses, the depth of subtrees to check increases, and isomorphic subtrees become more rare. Thus, iteration zero (where a subtree is a single node) dominates the similarity score. This phenomenon also poses challenges to the latter formulation by leading to the *diagonal dominance* problem [57], where a cellular graph is only identical to itself, not similar to any others. The population graph derived shows a substantial number of isolated nodes, representing patients dissimilar to any others. The community detection algorithm identifies five patient subgroups, none with distinct survival outcomes (see Fig. 17.d). These results suggest that the stringent comparison employed in the conventional WL subtree kernel may hinder the discovery of important patterns (e.g., patient subgroups) in a cohort. Additionally, the feature space of the WL subtree kernel has an extremely high dimension (see Extended Data Fig. 9), making the interpretation (i.e., identification of characteristic subtrees) prohibitive.
5. **Graph embedding:** Lastly, we use the FEATHER [58] graph embedding method to generate a low-dimensional vector for each cellular graph (i.e., patient). The inter-patient similarity is determined by the cosine similarities of two corresponding vectors. FEATHER models the spatial information and inter-cellular interactions using characteristic functions of node feature distribution. The resultant graph vectors are not explainable, and thus no explicit characteristic patterns can be identified. The population graph (Fig. 17.e) reveals clearer clusters compared with those using pairwise proximity of cell phenotypes (Fig. 17.b) and WL subtree kernel (Fig. 17.d). The community detection algorithm identifies five patient subgroups in the population graph, but none of these subgroups show distinct survival outcomes.

**Fig. 17:**
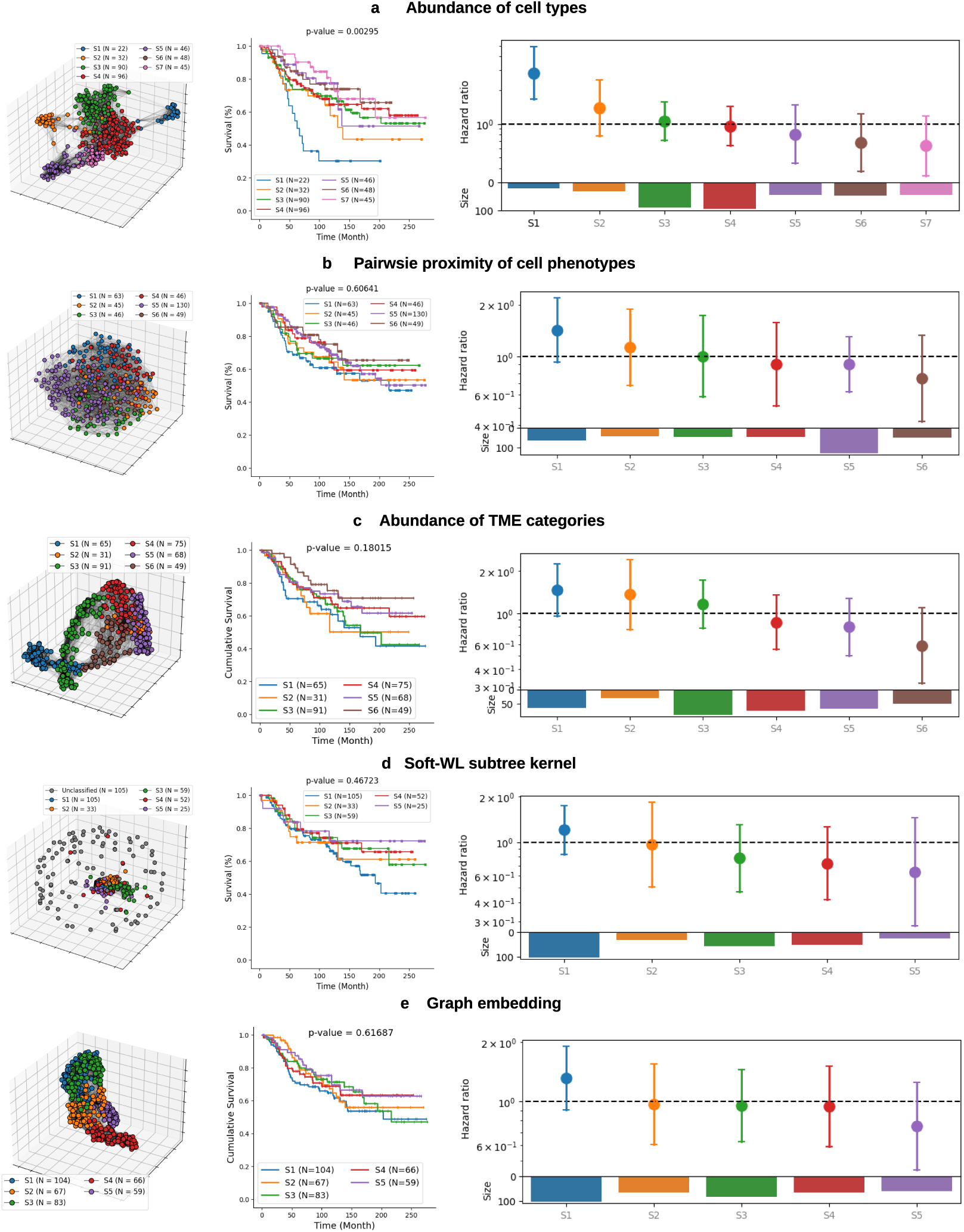
Results of comparative methods on the discovery set. **(a)** Risk stratification results from characterizing patients by the abundance of cell types on the discovery set. The first row shows the population graph with nodes painted by patient subgroup labels, the second row shows K-M survival plots of patient subgroups, and the third row shows hazard ratios estimated by the Cox proportional model. **(b)** Risk stratification results from characterizing patients by the pairwise distance of cell types on the discovery set. **(c)** Risk stratification results from characterizing patients by the abundance of four TME categories on the discovery set. **(d)** Risk stratification results of WL subtree kernel on the discovery set. **(e)** Risk stratification results of graph embedding method on the discovery set.

## 3 Discussion

The TME represents a fundamental piece in the study of cancer biology, and several recent studies indicate its promise for discovering biomarkers for prognosis and even treatment resistance [3, 4, 19, 20, 59–61]. However, the number of broadly applicable biomarkers from pathology that provide strong prognosis information, while generalizing to different studies, remains limited. Additionally, while some useful metrics, such as tumor-infiltrating lymphocytes (TILs) score [62, 63], tumor-immune mixing score [15], presence of tertiary lymphatic structure (TLS) [54, 64], among others, have been derived from pathology using traditional methods of segmentation tools (e.g. [65]) and spatial statistics [15], driven mostly by hypotheses relying on domain expertise, the discovery of data-driven biomarkers for prognosis has remained unexplored.

Modern deep learning methods, including graph neural networks, constitute an appealing alternative for data-driven biomarkers for prognosis [21, 22]. Yet, their employment in real-world clinical settings remains constrained by the lack of rigorous external validation [35, 36] and limited explainability [25, 26, 28, 29, 32, 33]. Instead, this study introduces a data-driven methodology centered on breast cancer, that relies on two simple observations: *i)* representative phenotypic patterns of the TME can be adaptively learned by studying the underlying patient-specific cellular graph constructed from multiplexed tissue image data, and *ii)* the relative abundance of such patterns in different patients can be employed to provide a measure of their similarity, from which a population-level graph can be constructed. This bi-level process allows us to obtain – automatically and in an unsupervised manner – different patient subgroups, with potentially distinct prognoses, either better or worse. Compared with other data-driven methods, such as graph-based deep learning, a significant strength of our methodology is the complete transparency of the features that provide the risk stratification and characterize better and worse survival. These features, dubbed *TME patterns*, represent similar subtrees in the TME, each corresponding to a local cellular neighborhood with unique cellular composition and spatial organization. As a result, we can easily unveil the most over-expressed structures that characterize better (and worse) survival, which elucidates the underlying associations between TME patterns and patients’ prognosis. Relying on more complex models, such as those based on graph neural networks, would have made it significantly more challenging – if not altogether impossible – to characterize such biomarkers clearly and easily.

Another contribution of this work is the formalization of the Soft-WL subtree kernel, a novel relaxation of the WL subtree kernel [43]. This kernel method efficiently captures and compares local structures in complex cellular graphs with numerous cells and various phenotypes. It over-performs other simpler methods, such as simply quantifying the abundance of different types of cells, by considering their spatial co-localization and interactions. On the other hand, it alleviates the diagonal dominance issue [57] of the conventional WL subtree kernel, providing a smoother comparison of subtrees. A limitation of this method is the over-smoothing phenomenon [66, 67] that is shared by many other graph learning methods. As the graph convolution progresses and subtree depth increases, subtrees tend to become increasingly similar to each other, preventing us from capturing higher-order structures in the graphs. We believe that BiGraph could be improved further, for example by replacing the Soft-WL subtree kernel with more advanced graph kernel methods.

In this study, we provide a data-driven risk stratification for a cohort of breast cancer patients, featuring seven subgroups of patients with varying prognostic patterns. This risk-stratification scheme is furthermore shown to be robust in both an inner validation set (a subset of patients randomly selected from the original cohort) and two external validation sets (independent datasets collected with different acquisition protocols and antigen panels). Patients in all validation sets studied (internal and externals) mirror the prognosis of those with similar TMEs in the discovery set. Furthermore, and importantly, by comparing this new risk stratification system with the standard clinical subtyping system based on the expression of HR and HER2, we found that our system provides complementary information that enhances the risk stratification capacity of the conventional guidelines.

By analyzing the TME patterns that characterize each patient subgroup, we observe that HER2^+^ tumor niches are strongly associated with worse survival, higher tumor grade, and clinically defined HER2+ subtype. These findings are validated in independent, external validation cohorts. As shown in many previous works, HER2 is an epidermal growth factor receptor, and its over-expression has been associated with a more aggressive breast cancer phenotype with decreased survival [68–71]. Our study is consistent with this observation, but it is found in an entirely data-driven manner. On the other hand, we found that tumor niches expressing high levels of cytokeratins and CXCL12 are associated with better outcomes, lower tumor grade, and the clinically defined HR+/HER2- subtype. This finding is particularly noteworthy as the prognostic impact of the interplay between these proteins, especially in breast cancer, is underexplored. While some literature discusses their individual impacts, controversial evidence on their prognostic impact exists: some demonstrate its association with the down-regulation of genes linked to metastasis inhibition, promoting invasion and migration of breast cancer cells [72, 73], while others suggest that higher expression of CXCL12 is associated with better survival, suppressing tumor growth and invasion in both breast [74] and pancreatic cancer [75]. The prognostic impact of cytokeratin is even more complicated due to the diverse range of subtypes. CK5/6, CK14, and CK20 are shown to be positively associated with high tumor grade of breast cancer [76, 77], and CK17 is also linked to poor clinical outcomes in breast cancer patients [78]. Conversely, down-regulation of CK8/18 has been reported to promote breast cancer progression, implying a favorable prognostic impact of CK8/18 [79–81]. Our study provides a new insight: that the co-expression of high levels of cytokeratin and CXCL12 is associated with better breast cancer prognosis. Importantly, this observation is found in an automatic, data-driven manner. The intricate mechanism underlying this association warrants further investigation.

Our work also focuses on TBNC patients in particular, given the severity of the disease in this clinical subtype. In this context, we discovered a unique TME pattern characterized by the dense aggregation of B cells surrounded by CD4+ T cells, CD8+ T cells, macrophages, and stromal cells. This pattern closely resembles the well-known TLS structure [54], and we denote it as “TLS-like” niche. Although the TLS-like niche is not associated with prognosis across the entire discovery cohort (i.e. including all cancer subtypes), it is linked to better survival among TNBC patients. This association is validated in two external validation sets, including a randomized clinical trial [47]. In particular, patients receiving combined chemotherapy and immunotherapy presented a statistically significant difference in the abundance of TLS-like niches in patients achieving pathologically complete responses and those with residual disease. These results suggest that TLS-like niches hold both prognostic and predictive values in TNBC patients. TLS, defined as the organized aggregation of immune cells in tumor sites resembling secondary lymphoid organs [82], is generally associated with promoted immune response and favorable clinical outcomes in a wide spectrum of cancer types in literature [82–85]. However, its role in breast cancer is mixed due to the disease’s heterogeneity. TLS presence has been linked to more aggressive tumor characteristics in breast cancer, such as higher proliferation and HR negativity [86] as well as the activation of tumor angiogenesis and invasion in animal models [87]. On the other hand, TLS is associated with better survival in HER2+ breast cancer, yet linked with increased lymphovascular invasion in Her2 negative patients [88]. Yet, evidence in TNBC is more consistent, with most studies suggesting TLS is associated with promoted immune response and better outcomes [89–92]. Our findings align with these works and are particularly noteworthy since they are discovered automatically from data, without providing our methodology with explicit prior knowledge and assumptions.

## 4 Conclusion

BiGraph employs a bi-level graph learning scheme to systematically integrate information across different scales: from local cell distribution to patient populations. In doing so, it allows for data-driven and explainable risk stratification and biomarker discovery even in small and heterogeneous cohorts, representing the potential to enhance our understanding of cancers and the development of personalized care. Naturally, this work has some limitations. The experiments and analyses presented in this work are limited to breast cancer, although the methodology is in principle applicable to other cancers. Due to the high cost of multiplexed technology, the sample sizes in this study are small, preventing us from making stronger statistical claims. Future work will focus on applying BiGraph to larger cohorts, and other disease types. On the methodological side, the Soft-WL subtree kernel proposed in this work captures small local structures (within approximately thirty microns) in the TME, sharing the oversmoothing obstacle common to many other graph-based algorithms. Moreover, although the soft-WL subtree kernel provides an effective heuristic relaxation of the classical WL subtree kernel, its broader capabilities and limitations require further theoretical investigation.

## 5 Methods

This section introduces the details of the methodology developed and applied in our study.

### 5.1 Preliminaries

We first present some key notation and definitions concerning graphs to facilitate the introduction of other methodologies.

A *graph* 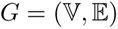 is defined by a tuple of a set of nodes 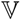 and a set of edges 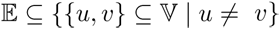. The set of nodes and the set of edges for a graph *G* can be written as *V* (*G*) and *E*(*G*), respectively. For each node *v*, the set of nodes with an edge connected to a node *v* is defined as its neighborhood, denoted as *N* (*v*). The structure of a graph can be fully characterized by its adjacency matrix, denoted as *A*. An adjacency matrix with only binary entries represents a binary graph. Conversely, an adjacency matrix with continuous scalar entries represents a weighted graph, where *A_uv_* indicates the weight of the edge connecting nodes *u* and *v*. Two graphs, *G*_1_ and *G*_2_, are deemed *isomorphic* if there exist a bijection *φ* : *V* (*G*_1_) → *V* (*G*_2_), such that (*φ*(*u*)*, φ*(*v*)) ∈ *E*(*G*_2_) for any (*u, v*) ∈ *E*(*G*_1_).

For a graph *G*, a *subtree* can be derived given a root node *v*, with the neighbors *N* (*v*) serving as its children. This subtree can be expanded by iteratively adding the neighbors of the children, akin to increasing the depth of a subtree. A subtree rooted at *v* with a depth of *h* is denoted *T ^h^*(*G, v*) (see Fig. 18). Two subtrees are considered *isomorphic* if they exhibit identical node sets at every level.

**Fig. 18:**
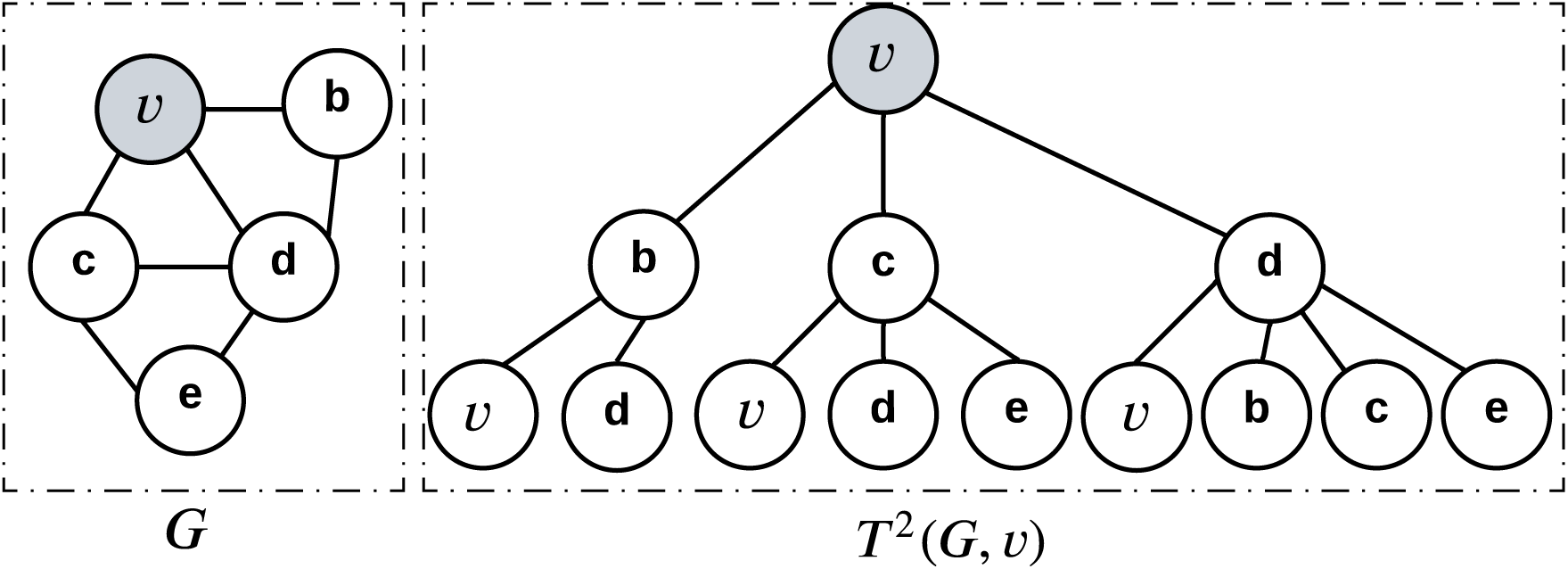
An illustration of a subtree rooted at node *v* (shaded) with a depth of 2.

### 5.2 Construction of cellular graphs from TMA slides

The construction of the cellular graph involves representing the tumor microenvironment (TME) based on spatial cell coordinates and cell phenotypes. Each node in the cellular graph represents a single cell. The cell phenotype serves as the label of each corresponding node in the graph, determined by clustering multi-channel protein expression data. In this study, we use the published phenotype results of the discovery set [16] and the validation set [17], respectively. Edges in the cellular graph represent inter-cellular connections. Previous approaches typically employed a fixed distance threshold (*d*_0_) to create a binary cellular graph, where two cells are connected only if the distance between them is smaller than *d*_0_. However, there is no consensus on the appropriate value for *d*_0_ and various choices have been employed, such as 4 *µ*m [30], 8 *µ*m [16], and 32 *µ*m [15]. Additionally, some of the works count the distance between cell centroids, while others consider the minimal distance. To provide a more comprehensive representation of inter-cellular interaction, our approach constructs a weighted cellular graph, wherein all possible pairs of cells are connected and the strength of the connection decreases with increasing inter-cellular distance. The edge weight (i.e., *w_uv_*) is calculated via a Gaussian kernel given by

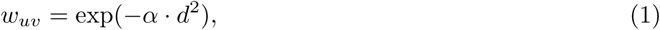

where *d* represents the spatial (Euclidean) distance between centers of cells *u* and *v* in micrometers (*µ*m), and *α* is a parameter set to 0.01 to calibrate the edge weights. The choice of *α* determines how fast these weights decrease with distance. As a concrete example, cells with a distance smaller than 3 *µ*m have strong connections with edge weights higher than 0.9, while cells with a distance larger than 22 *µ*m have weak connections with edge weights smaller than 0.01. In cases where a patient has multiple tissue microarray (TMA) cores, a separate cellular graph is constructed based on each core image, and then these individual graphs are combined as disconnected components to form a larger cellular graph representing the entire TME for that patient. This approach makes it possible to accommodate patients with multiple images while preserving the individual characteristics of each cellular graph and maintaining the entire methodology unaltered.

### 5.3 Soft-WL subtree Kernel

In this section, we delve into the Soft-WL subtree kernel employed in our work, which is a relaxation of the well-known Weisfeiler-Lehman (WL) kernel [43]. The WL subtree kernel, as detailed in Section 5.8, relies on isomorphic subtrees as a similarity measure across graphs, and is only applicable to binary graphs. The Soft-WL subtree kernel is designed to handle weighted and complete cellular graphs and provides a smoother comparison between subtrees. Fig. 19 shows a toy example of the Soft-WL subtree kernel. It comprises three main steps: (1) generation of subtrees (see Fig. 19.a); (2) identification of TME patterns by clustering subtrees (see Fig. 19.b); and (3) measurement of inter-patient similarity based on the abundance of subtrees (see Fig. 19.c). We detail each of them below.

**Fig. 19:**
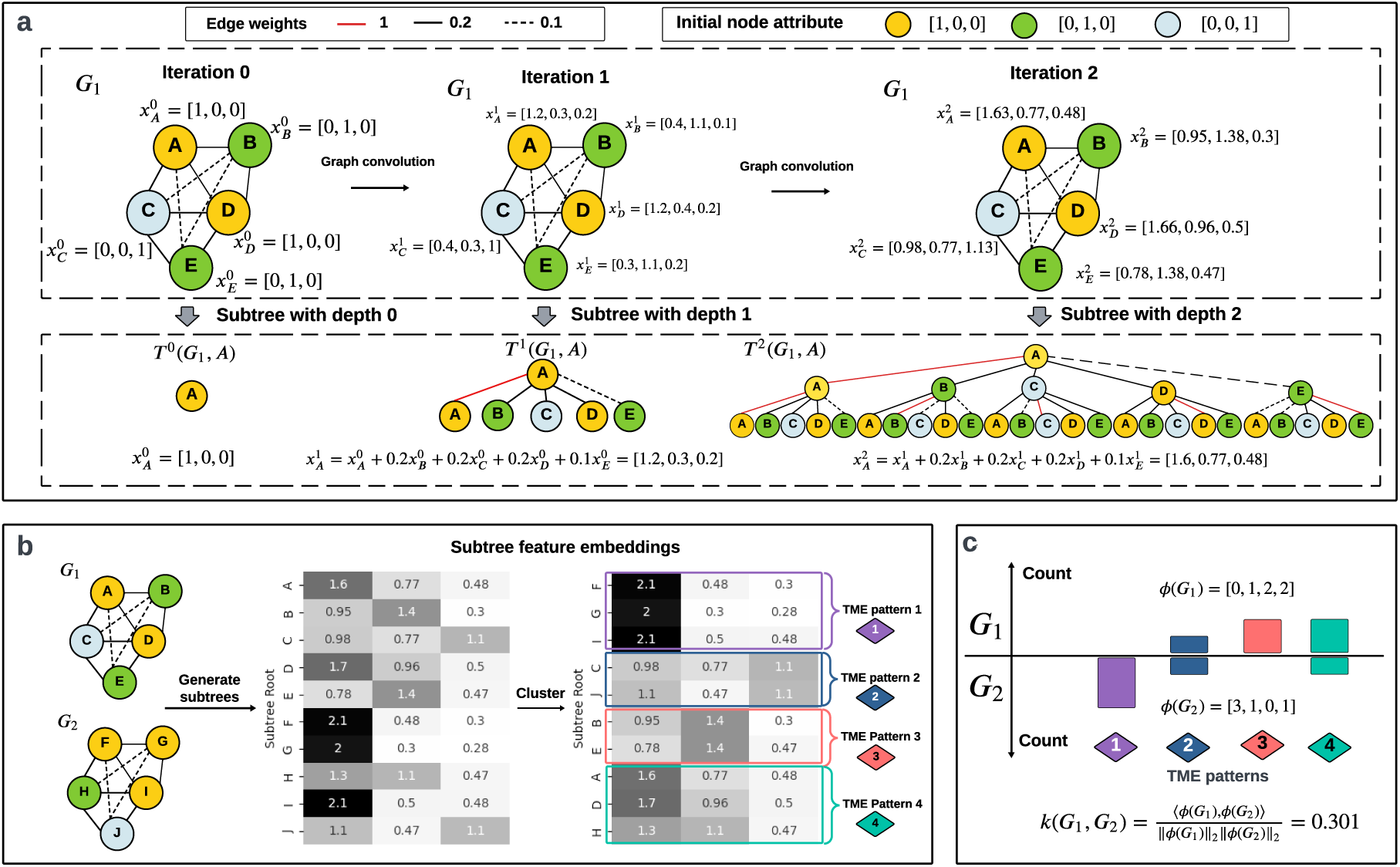
A toy example of the Soft-WL subtree kernel applied to two small cellular graphs. **(a)** The process of generating subtrees from the small cellular graph *G*_1_ with five nodes is illustrated. Nodes are initially assigned attributes representing the cell type (denoted by node color), and these attributes are iteratively updated through a graph convolution process. At each iteration, the updated node attribute serves as the feature embedding of a subtree rooted at that node with a designated depth. Subtrees rooted at node A with depths 0, 1, and 2 are displayed. **(b)** Clustering of the ten subtrees generated from two cellular graphs, *G*_1_ and *G*_2_, each rooted at a single node with a depth of 2. The resulting four clusters correspond to four TME patterns. **c** The similarity score between *G*_1_ and *G*_2_, denoted as *k*(*G*_1_*, G*_2_), determined by the normalized inner product of their histograms of the four TME patterns.

#### 5.3.1 Generation of subtrees

Given that the cellular graph constructed is weighted and complete, with each cell connected with all others with varying weights, the resulting subtrees are typically large and weighted, leading to a rare occurrence of isomorphic subtrees. Instead, a feature embedding is computed for each subtree through a graph convolution process. Formally, we consider a cellular graph 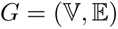 with a weighted adjacency matrix *A*. Noticeably, self-loops are added to each node, and thus the diagonal entries of *A* are all 1. Each node, *v*, is initially assigned an attribute, which is the one-hot encoding of its cell phenotype, denoted as 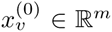, where *m* is the number of unique cell phenotypes. The node attribute is then iteratively updated as follows:

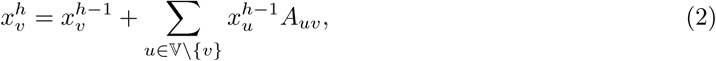

where *h* is the iteration index. Intuitively, this graph convolution process enables the updated node feature to incorporate not only its phenotype information but also that of its neighbors. Thus, the updated node feature, 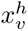, is considered the feature embedding of the subtree rooted at *v* with a depth of *h*, denoted as *T ^h^*(*G, v*). In this way, this feature embedding encodes the structure of a local cellular neighborhood surrounding *v*. Note that the cellular neighborhood does not have a clear boundary, but one can decide on a boundary by applying a threshold *τ* to the impact of neighborhood nodes on this feature embedding. To be more specific, Equation (2) can be rewritten as

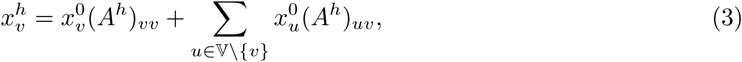

where *A^h^* denotes the *h*^th^ power of the adjacency matrix *A*, and 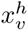 is 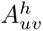 denotes the influence of a node *u* to 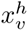. Then, any node *u* with (*A^h^*)*_uv_ > τ* is considered part of the cellular neighborhood. This hyper-parameter, *τ*, is user-defined, and we set it as 0.01 in the experiments. In this way, *N* subtrees can be generated from a cellular graph with *N* cells, each rooted at a single cell, and having a feature embedding derived from the graph convolution process above. The feature embedding of each subtree represents the structure of a local cellular neighborhood surrounding each cell, which is the root of the tree.

#### 5.3.2 Identification of TME patterns

Unlike the traditional WL kernel [43], which focuses on the distribution of subtrees as a similarity measure, the Soft-WL subtree kernel clusters similar subtrees *T ^h^*(*G, v*) based on their feature embeddings 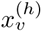. Moreover, instead of comparing subtrees embedding themselves, we will compare the distribution of *cell patterns* across patients as characterized by their embeddings.

To this end, we employ a clustering algorithm across subtrees from the entire discovery set. Although the clustering method can be general, we employ the PhenoGraph algorithm [45] since it has been shown to be less sensitive to hyperparameters in the context of single-cell data. The official Python implementation from PhenoGraph (https://github.com/jacoblevine/PhenoGraph) is used. Each resulting cluster is considered a *TME pattern*, which thus characterizes similar subtrees (i.e. those having similar feature embeddings). Furthermore, each TME pattern is assigned a signature, denoted `sign`, given by the cluster centroid. More precisely, for each *i*^th^ out of *K* clusters,

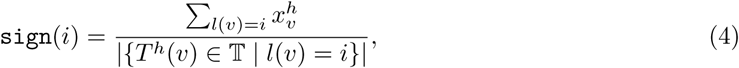

where 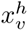 is the subtree feature embedding of node (cell) *v*, 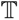 denotes the set of subtrees derived from the entire discovery set, and *l* : *V* → {1*,…, K*} denotes a function mapping a subtree to its assigned cluster identity.

#### 5.3.3 Inter-patient similarity measurement

The Soft-WL Subtree kernel calculates the inter-patient similarity score by comparing the abundance of these TME patterns across patients. Let *ϕ*(*G*_i_) be the histogram of TME patterns in the graph of each subject *G*_i_. This similarity is then given by

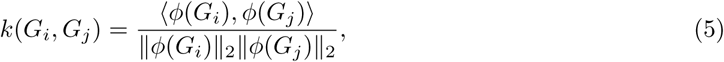

where ‖·‖_2_ denotes the *ℓ*_2_ (Euclidean) norm. The normalization ensures that the similarity score for any pairs of patients falls within the range of 0 to 1, with identical graphs achieving a maximal similarity of 1. The hyperparameter, *h*, controls the depth of subtrees and indirectly decides the size of the corresponding local cellular neighborhood. We set it as *h* = 2 in the experiments. As detailed in Section 5.3.1, the cellular neighborhood boundary is decided by the contributions of surrounding cells to the subtree embedding. The cellular neighborhoods contain a median of 16 cells, with a median radius (defined by the maximum distance between a subtree root and the cells within that cellular neighborhood) of 30 um.

### 5.4 Population graph and community detection

In the previous section, patient-specific graphs are constructed to characterize the local statistics of cell distribution in terms of recurrent patterns. In contrast, the population graph that we describe now is constructed to encode the similarities *across* patients. Each node of the population graph represents a patient, and every patient is connected to potentially all other patients through edges with scalar weights. Such weights represent the similarity between any two patients measured by the Soft-WL subtree method, given by Equation (5). Thus, the patient graph can be represented by the *Kernel matrix K* ∈ ℝ^N×N^ for the *N* patients {*G*_1_*, G*_2_*, …, G*_N_ } in the discovery set, and each entry *K_ij_* is given by *K_ij_* = *k*(*G_i_, G_j_*) (Equation (5)).

The population graph is preprocessed in the following two steps before proceeding to detect sub-communities within this graph (as per instructions in [45]): (1) identifying the *k*^⋆^-nearest neighbors (*k*^⋆^-NN); (2) replacing the edge weight between two patients by the intersection-over-union of their respective *k*^⋆^-NN sets. The Louvain community detection method [46] (Python package “networkx” [93]) is then employed to detect communities within the population graph, where each community represents a patient subgroup characterized by high intra-group similarities in their TMEs. We set the hyperparameter *k*^⋆^ = 30 in the experiments. Although this hyperparameter impacts the partition granularity, the patient subgrouping results show substantial consistency across various choices of *k*^⋆^ (see Extended Data Fig. 2).

### 5.5 Identification of characteristic patterns

The pattern histogram is normalized on a per-patient basis by dividing the occurrence of each pattern by the sum of occurrences of all patterns within that patient, yielding the proportion of each TME pattern in individual patients (see Extended Data Fig. 3). Consequently, the Hodges–Lehmann statistic [53] is employed to compare the distribution of a specific TME pattern’s proportions among patients inside and outside a patient subgroup (Extended Data Fig.4). A higher Hodges–Lehmann statistic indicates the increased expression of the studied pattern in the patient subgroup. A TME pattern is deemed “characteristic” if its Hodges–Lehmann statistic surpasses 50% of the maximum value within that patient subgroup, as highlighted in colored boxplots in Extended Data Fig. 4.

### 5.6 Prognosis analysis

The Kaplan-Meier (K-M) estimator [49] is used to generate survival plots for each patient subgroup. For the discovery set and inner validation set, the disease-specific survivals are considered. On the other hand, the overall survival is considered for the external validation set, since the disease-specific survival data is not available. The multivariate log-rank test [50] is used to compare their survivals. A Cox proportional hazard regression model [51] is used to calculate hazard ratios for each patient subgroup, with patients outside each patient subgroup serving as the baseline. In assessing the prognostic impact of a specific TME pattern, patients are stratified into positive and negative groups based on the expression of the target TME pattern, where a proportion of 1% is used as the threshold for categorization. The survival outcomes of positive and negative patients are compared using the pairwise log-rank test. All of these prognosis-related statistical methods are conducted with the Python implementation from “lifelines” [94].

### 5.7 Validation and generalization

The robustness and generalization of the major findings in the discovery set are rigorously assessed in both an inner validation set and an external validation set. The validation process encompasses mapping at the cellular, subtree, and patient levels. We detail each of them in the following.

#### 5.7.1 Cell phenotyping system mapping

Cell phenotype mapping is only needed for the external validation set since it employs a different cell phenotyping system from that in the discovery set. Moreover, the external validation set also employs a different antigen marker panel, with approximately 50% of the antigens shared between two sets. Each cell, irrespective of its origin in the discovery set or the external validation set, is characterized by a continuous feature vector representing the expression profiling of the *shared* antigens. For a cell from the external validation, mapping is achieved by assigning it to the phenotypically closest cell phenotype (i.e., a cluster of cells) defined within the discovery set. The phenotypic distance between a cell and a cell cluster is given by the Euclidean distance between the cell’s feature vector and the cluster centroid.

#### 5.7.2 TME pattern mapping

For each new patient in the validation set with its corresponding cellular graph, subtree generation, and feature computation are carried out in an analogous manner to that described in Section 5.3.1. Instead of re-conducting clustering on these subtrees from the validation set to find new patterns, they are mapped to the closest TME pattern (i.e., a cluster of subtrees) previously defined within the discovery set (i.e. allowing us to assign a new patient to a predefined subgroup). The distance between a subtree and a TME pattern is given by the Euclidean distance between the subtree’s feature vector and the pattern’s signature (i.e. cluster centroid).

#### 5.7.3 Patient subgroup mapping

Once TME patterns are mapped, the similarity between patients across sets can be calculated using Equation (5). Given a new patient in the validation set, its top three most similar patients in the discovery set are identified. The new patient is then mapped to the patient subgroup via weighted majority voting among the three nearest neighbors. In this way, a corresponding patient subgroup, denoted as Si^′^, can be identified for every group Si, both in the inner validation and external validation sets. Survival analysis is performed on them to discern whether a mapped patient subgroup mirrors the survival outcome of its phenotypically similar patient subgroup defined within the discovery set.

### 5.8 Comparative methods

This section outlines alternative methods for measuring inter-patient similarity scores considering Equation (5) as a general formula, but where *ϕ*(*G*) varies for each method.

- Abundance of cell types: In this method, *ϕ*(*G*) represents the histogram of cell phenotypes within a patient (i.e., corresponding cellular graphs).
- Pairwise proximity of cell phenotypes: Given a dataset with *n* unique cell phenotypes, *ϕ*(*G*) is a feature vector with 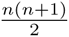 entries, each corresponding to the average proximity score between a pair of cell phenotypes given by:

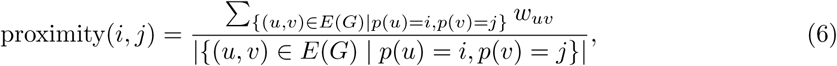

where *i* and *j* are two cell phenotypes (where *i* may equal to *j*), *E*(*G*) denotes the set of edges, *w_uv_* is the proximity score between cell *u* and *v*, akin to the edge weight given by Equation (1), and *p* is a function mapping each node (i.e., cell) to the corresponding cell phenotype.

- WL-subtree kernel [43]: This kernel measures the similarity between two graphs by counting the occurrence of isomorphic subtrees in them. Notably, the WL subtree kernel only takes binary graphs; therefore: weighted cellular graphs are converted to binary graphs by applying a threshold of 0.01 to the edge weights. This threshold connects two cells if the distance between their centroids is less than 21 um. The identification of isomorphic subtrees involves a *color refinement* procedure, where node colors are iteratively updated based on their neighbors. To be more specific, each node is initially assigned a color based on its cell phenotype, denoted as *σ*^0^(*v*). The node color can be either a discrete alphabet or an integer. The node color is subsequently updated as follows:

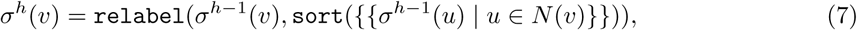

where *σ^h^*(*v*) is the node color at iteration *h*, and *N* (*v*) represents the neighbors of *v*. The function `sort` collects and sorts the multiset of colors of *N* (*v*), and the function `relabel` maps the pair (*σ^h^*^−1^(*v*), sort(*σ^h^*^−1^(*u*) | *u ∊ N* (*v*))) to a unique color not used in previous iterations. Two nodes with the same color after iteration *h* imply the isomorphism of the two subtrees rooted at these nodes with depth *h*. Notably, the color refinement is conducted universally for all the graphs in the dataset rather than individually, ensuring a consistence color system across all graphs. In this case, the feature *ϕ*(*G*) represents the histogram of unique colors, and two slightly different formulations are considered. The first formulation considers the occurrence of colors from all previous iterations, given by

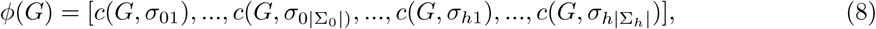

where *σ_hi_* is a unique color appearing at the *h*-th iteration of color refinement, Σ*_h_* = {*σ_h_*_1_*, …, σ_h_*_|Σ_*_h_*_|_} denotes the set of unique colors at the *h*-th iteration, and *c*(*G, σ_hi_*) denotes the occurrence of color *σ_hi_* in *G*. The second formulation only considers the colors from the last iteration of color refinement, given by

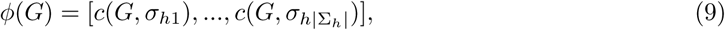

The hyperparameter *h* controls the depth of subtrees, similar to its counterpart in the Soft-WL subtree kernel. We set it to 2 in the experiments, consistent with the setting used for the Soft-WL subtree kernel. The WL subtree kernel is implemented with the Python package “Grakel” [95].

## Acknowledgement

This study is supported by the National Institutes of Health grant R01CA138264.

## Data availability

The multiplexed image data and corresponding clinical data used in this work are publically available at https://doi.org/10.5281/zenodo.5850952 (Danenberg et al [16]), https://doi.org/10.5281/zenodo.3518284 (Jackson et al [17]), and https://doi.org/10.5281/zenodo.7990870 (Wang et al [47]).

## Code availability

The code used for implementing the BiGraph method and all the analysis in this paper is available at https://github.com/Sulam-Group/BiGraph4TME.git

## Declaration of generative AI and AI-assisted technologies in the writing process

During the preparation of this work the authors used the large language model (ChatGPT 3.5) in order to improve conciseness. After using this tool/service, the authors reviewed and edited the content as needed and take(s) full responsibility for the content of the publication.

---

**Extended Data Fig. 1:**
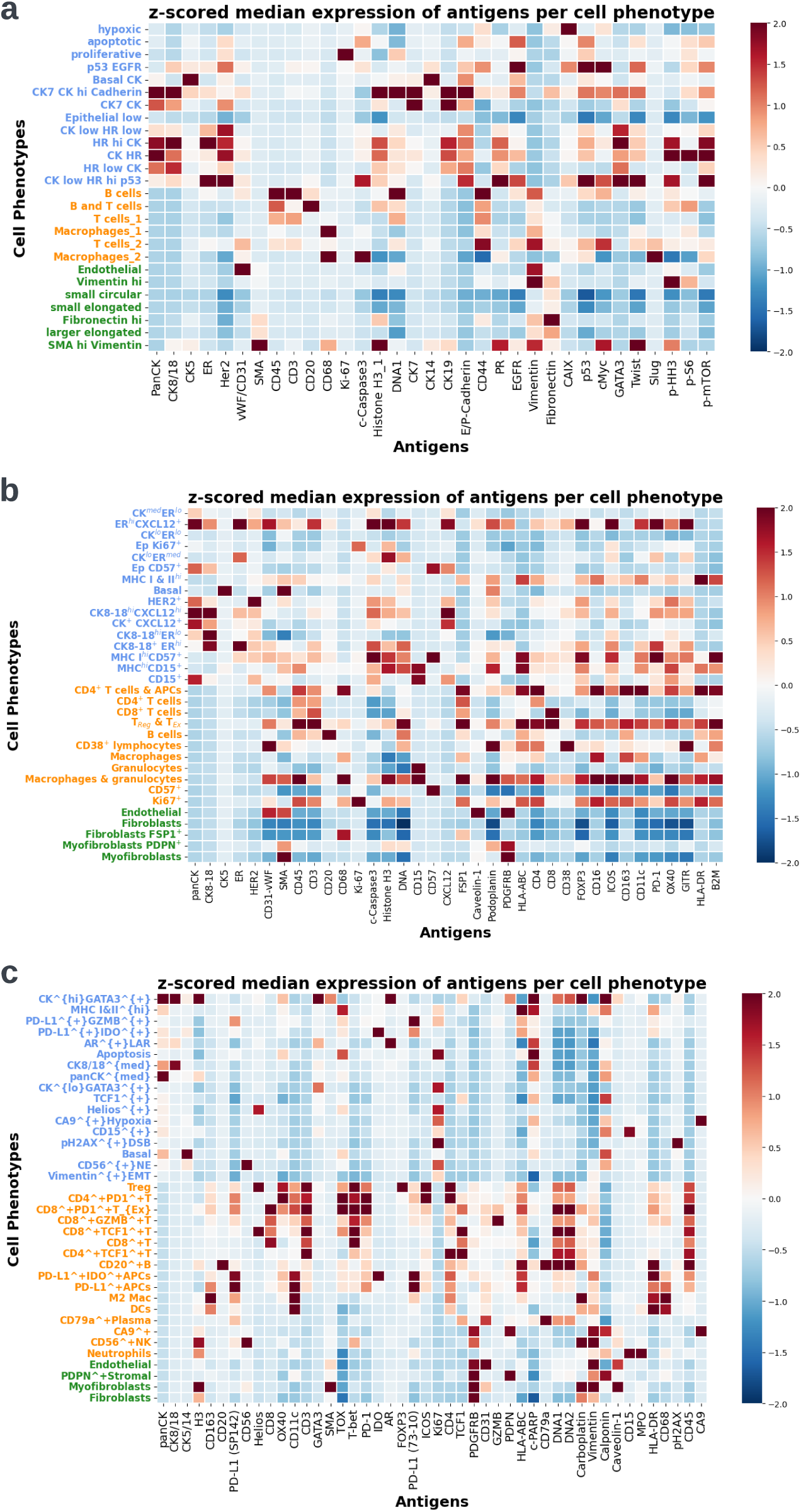
Cell phenotyping system of the datasets. **(a) - (c)** The median antigen profiling of different cell phenotypes in the discovery set curated by Danenberg et al. [16], external validation set-1 curated by Jackson et al. [17], and external validation set-2 curated by Wang et al. [47]. The y-axis label colors indicate the cell categories: blue: tumor cells; orange: immune cells; green: stromal cells.

**Extended Data Fig. 2:**
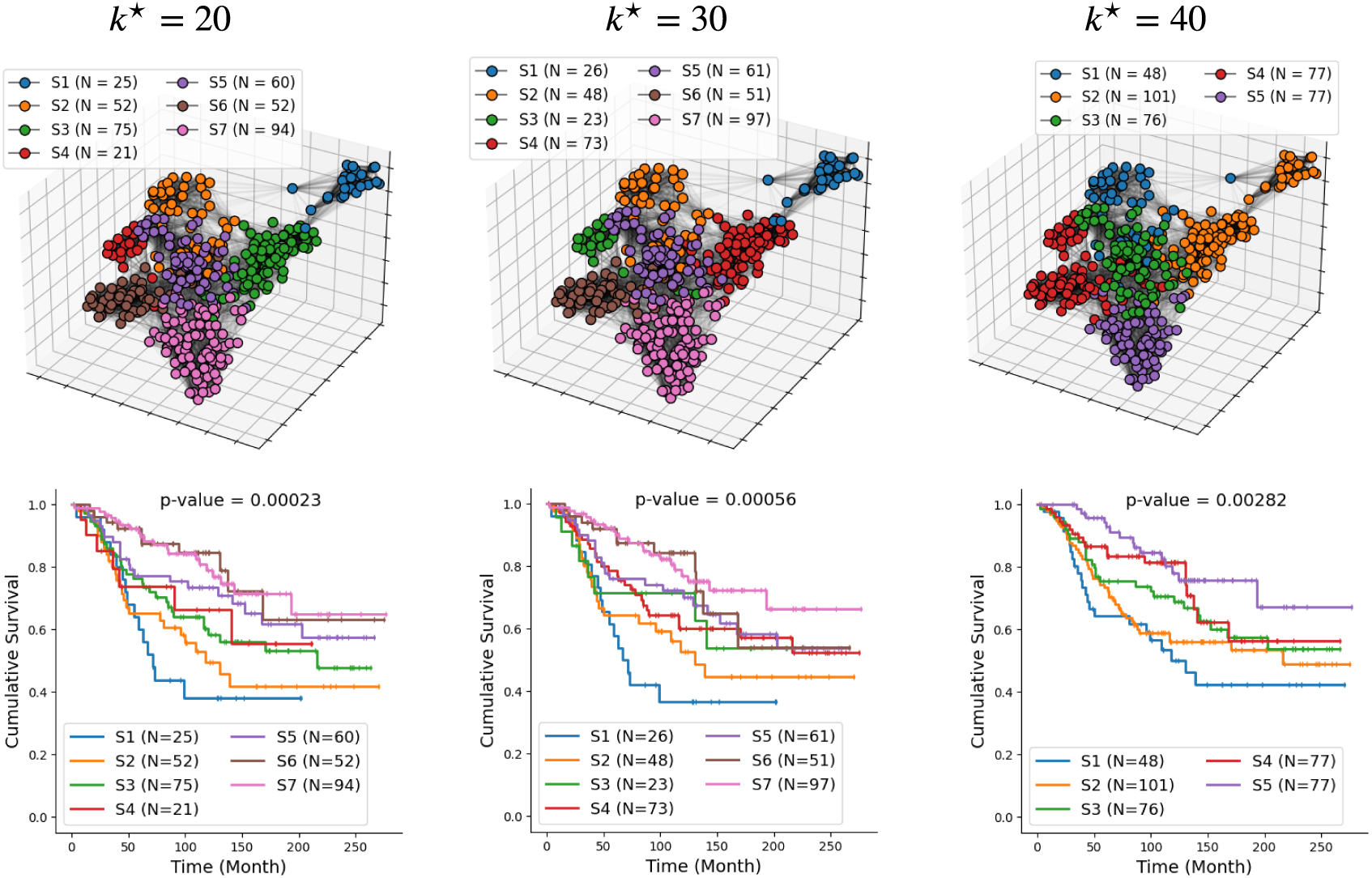
Detected patient subgroups with different choices of hyperparameter. *k*^⋆^. The first row displays population graphs with nodes painted by patient subgroup ids, and the second row shows the survival plots of patient subgroups. Multi-variate log-rank test is used to compare their survivals, and the p-values are shown on the titles.

**Extended Data Fig. 3:**
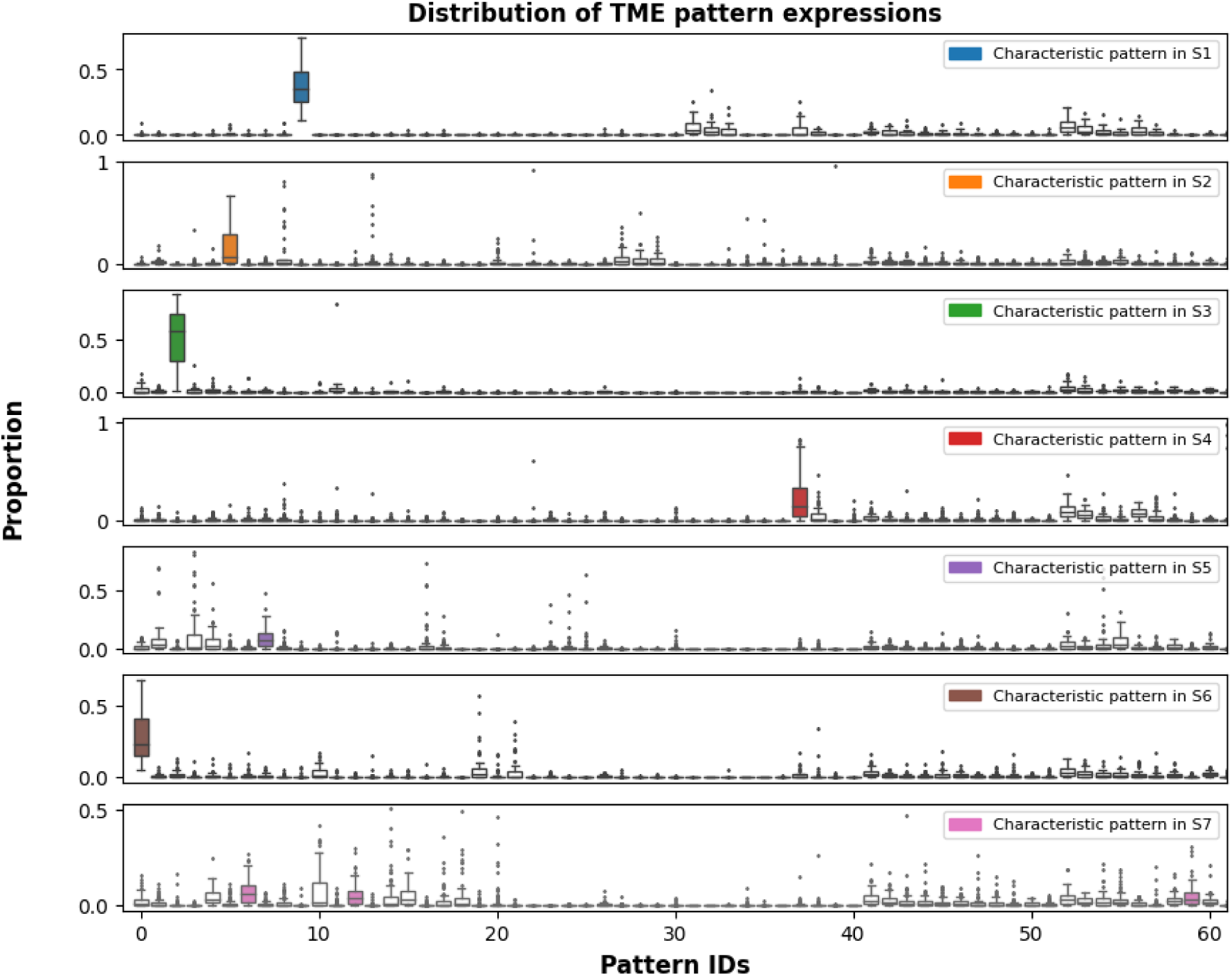
Distribution of TME pattern expressions in each of the seven patient subgroups. The proportion of each specific TME pattern within an individual patient is calculated by normalizing the corresponding TME pattern histogram. The distribution of these proportions within each patient subgroup is visually represented using box plots. Characteristic patterns are highlighted by colored box plots.

**Extended Data Fig. 4:**
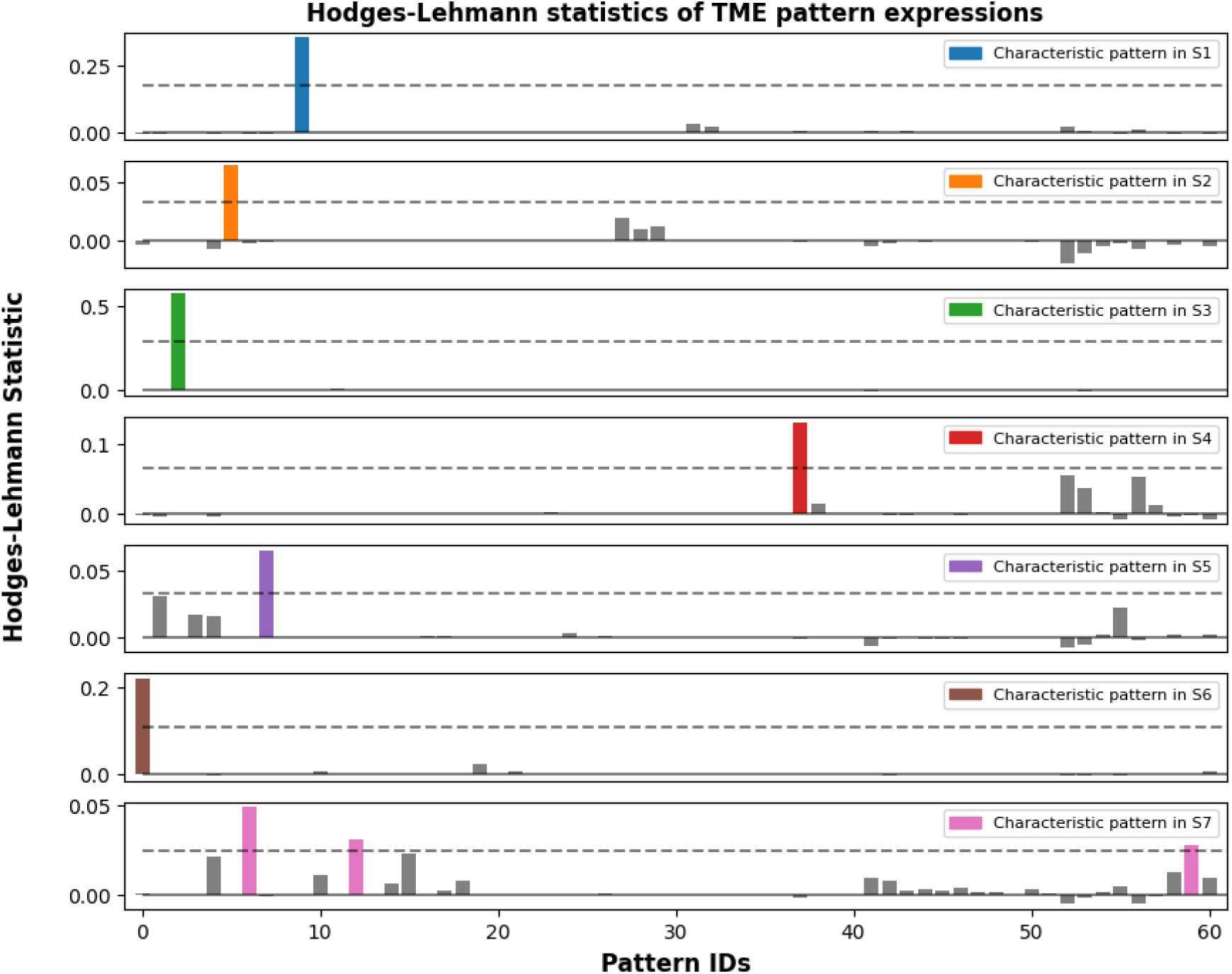
Hodge-lehmann statistics of TME pattern expressions in each of the seven patient subgroups. For each specific TME pattern within a designated patient subgroup, the distribution of its proportions among patients inside and outside that patient subgroup is compared using the Hodges–Lehmann statistic. A larger Hodges–Lehmann statistic infers an over-expression of that particular pattern in that patient subgroup. A TME pattern is deemed “characteristic” in a patient subgroup if the Hodges–Lehmann statistic exceeds 50% of the maximized value within that patient subgroup, highlighted by colored bar plots.

**Extended Data Fig. 5:**
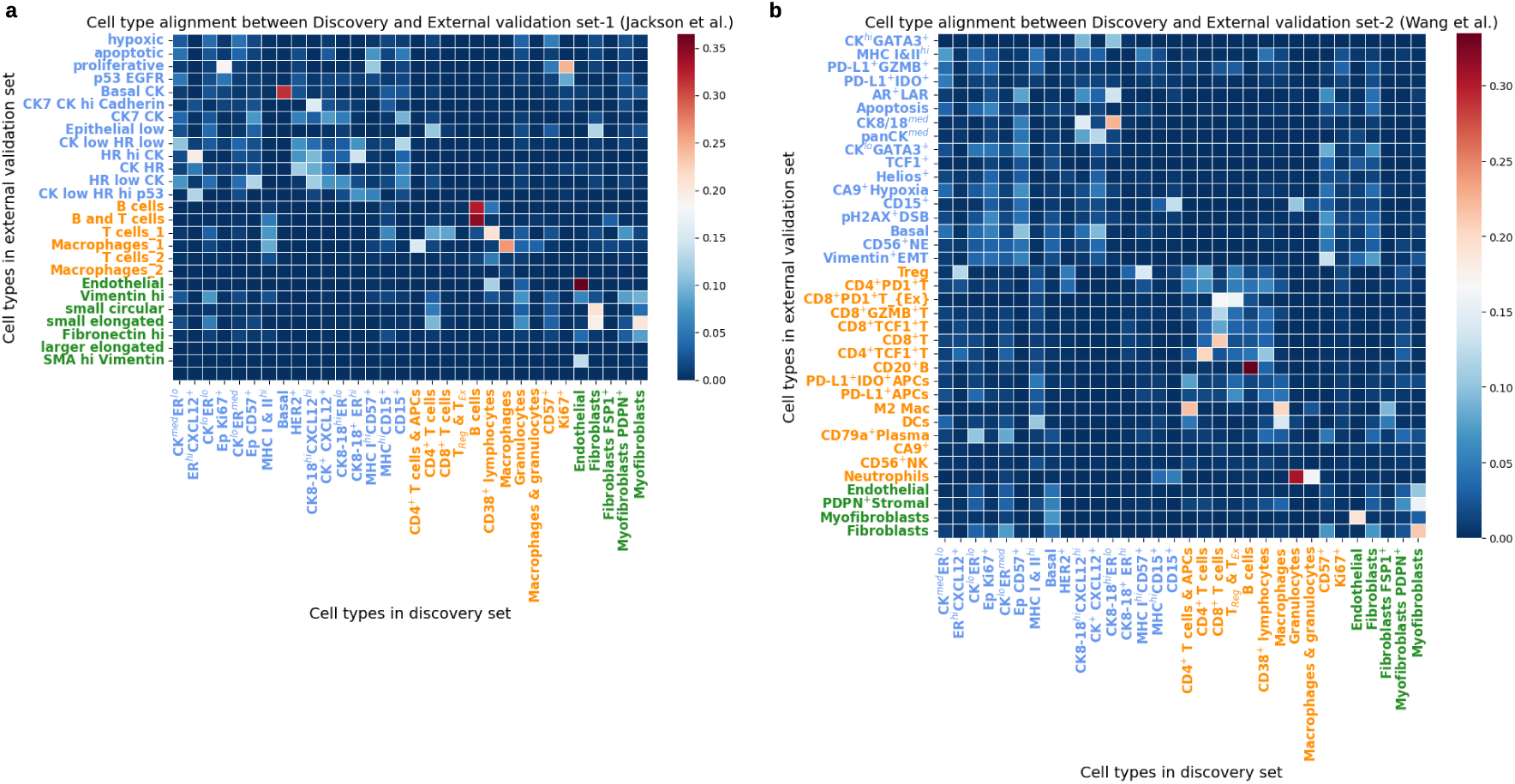
Cell phenotyping system alignment between discovery set and external validation set. Each cell within the external validation set is associated with two distinct phenotype labels: firstly, the original label as designated in the external validation set phenotyping system; and secondly, the label assigned within the phenotyping system of the discovery set, determined through cell phenotype mapping. The Intersection over Union (IoU) is computed for each pair of cell phenotypes derived from these two distinct phenotyping systems, and the results are visually represented in the heatmap. **(a)** The alignment between discovery set [16] and external validation set-1 [17]. **(b)** The alignment between discovery set [16] and external validation set-2 [47].

**Extended Data Fig. 6:**
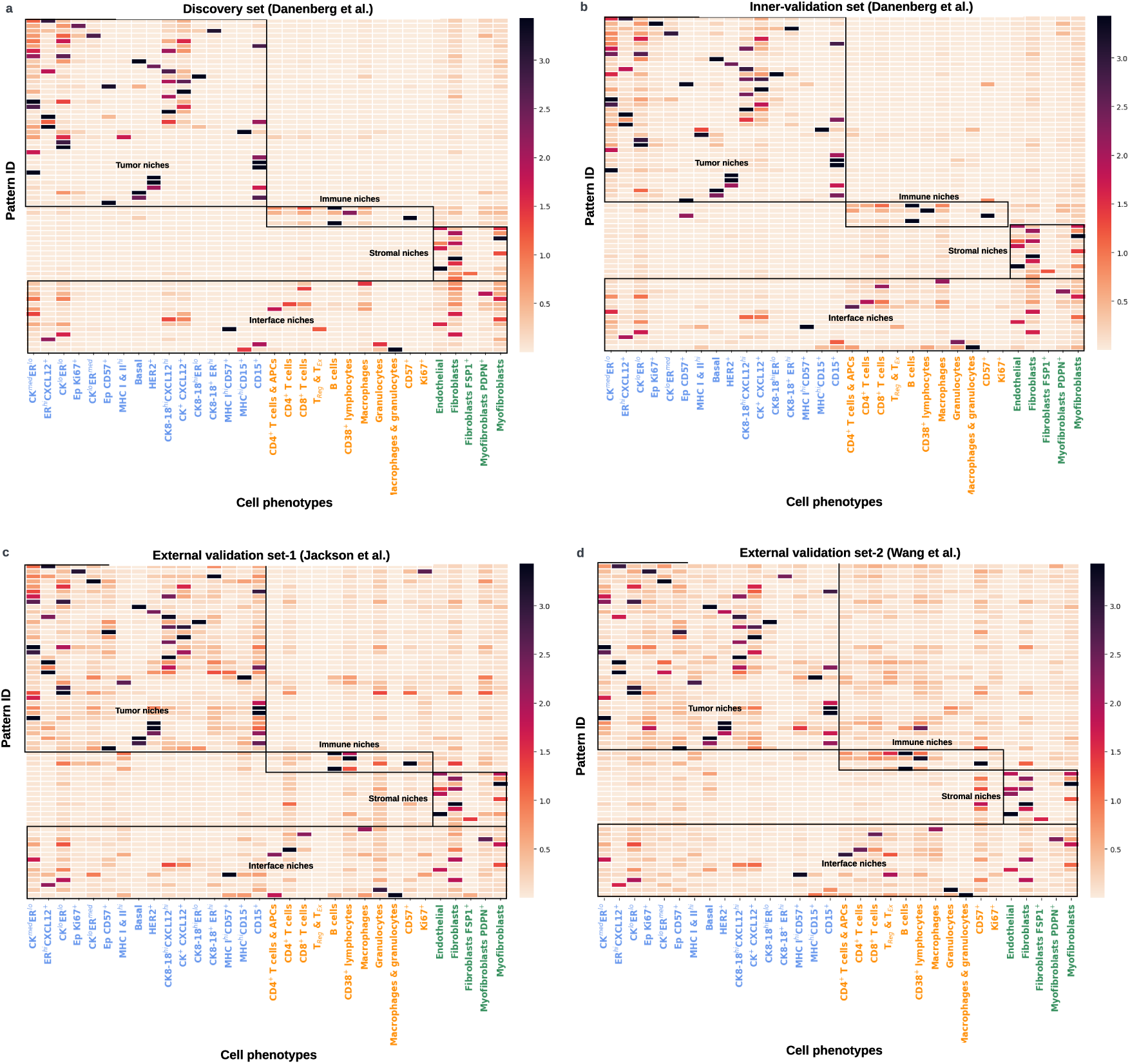
Signature maps of 66 TME patterns across different datasets. **(a)** The signature map (cluster centroids) of 66 TME patterns identified in the discovery set [16]. **(b)-(d)** In the validation sets, each unseen subtree is mapped to one of these 66 predefined TME patterns. The figures show the signature maps (cluster centroids) of the 66 TME patterns in the inner-validation set, external validation set-1 [17], and external validation set-2 [47].

**Extended Data Fig. 7:**
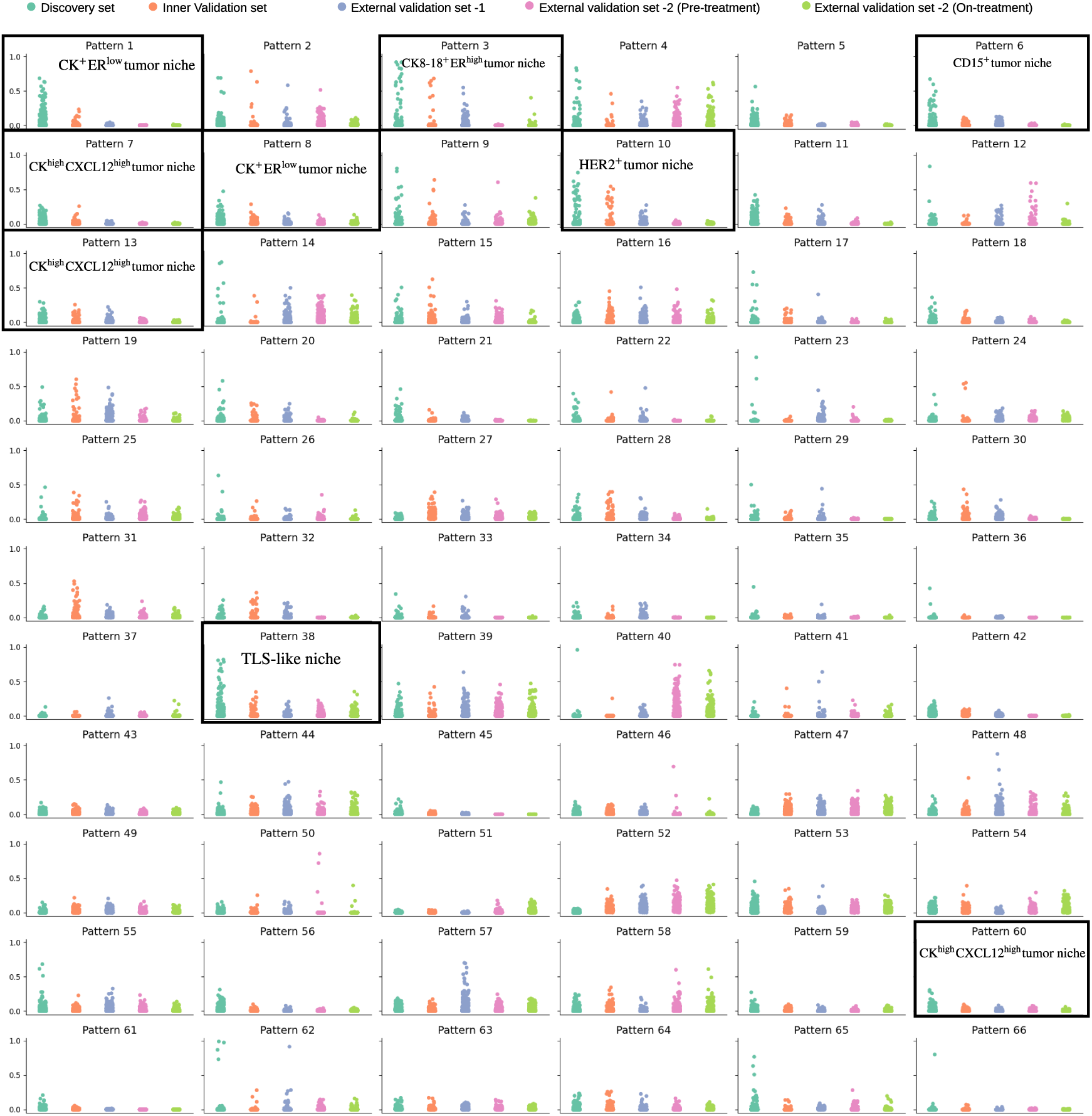
Distribution of 66 TME patterns across different datasets. Each panel presents the distribution of a specific TME pattern, as the title indicates. The x-axis represents the datasets, while the y-axis shows the proportion of the TME pattern. Each small circle corresponds to the proportion of a specific TME pattern in an individual patient. Different colors represent corresponding datasets: cyan for the discovery set, orange for the inner validation set, blue for external validation set-1, purple-red for external validation set-2 (pre-treatment samples), and lime for external validation set-2 (on-treatment samples).

**Extended Data Fig. 8:**
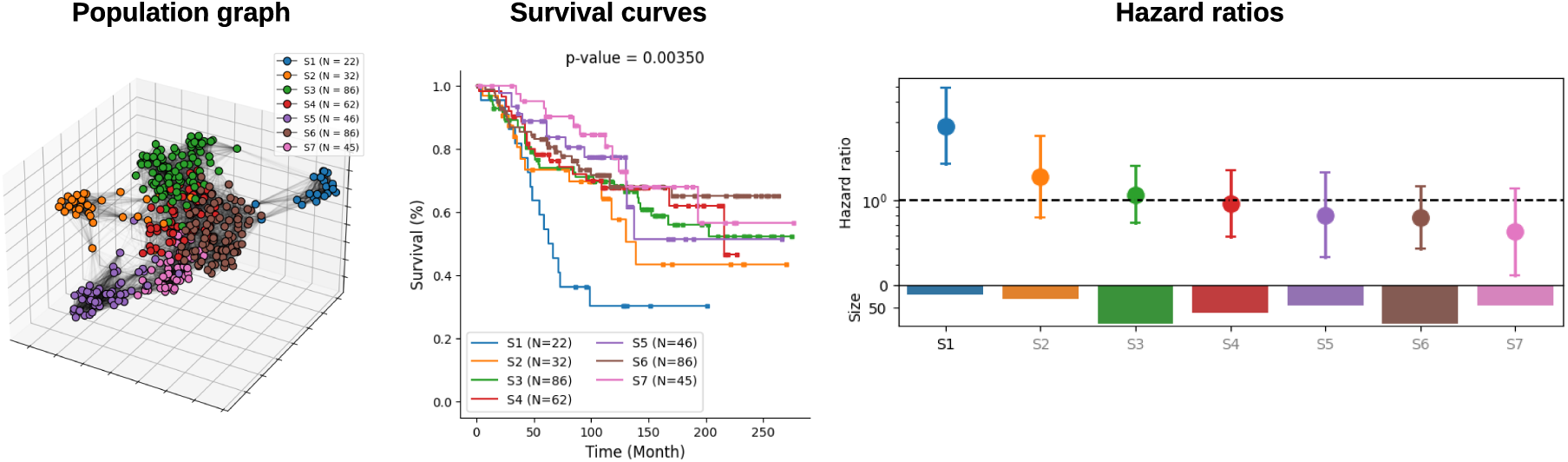
Risk stratification results from WL subtree kernel (accumulating similarities from all previous iterations). From left to right, the figure shows a population graph with nodes painted by patient subgroup labels; K-M survival plots of patient subgroups; and hazard ratios estimated by the Cox proportional model.

**Extended Data Fig. 9:**
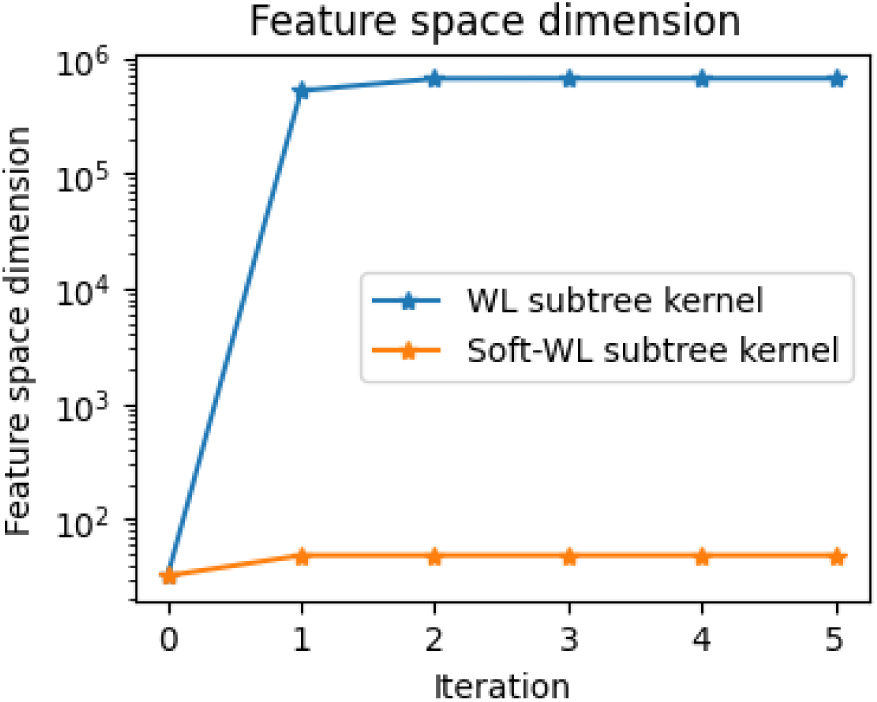
Feature space dimension of WL and Soft-WL subtree kernels. To the WL subtree kernel, the feature space dimension refers to the number of unique colors of each iteration of color refinement. To soft-WL subtree kernel, the feature space dimension refers refers to the number of TME patterns at each iteration of graph convolution.

